# Disentangling the Neural Correlates of Agency, Ownership and Multisensory Processing

**DOI:** 10.1101/2022.08.15.504036

**Authors:** Amir Harduf, Ariel Shaked, Adi Ulmer Yaniv, Roy Salomon

## Abstract

The experience of the self as an embodied agent in the world is an essential aspect of human consciousness. This experience arises from the feeling of control over one’s bodily actions, termed the Sense of Agency (SoA), and the feeling that the body belongs to the self, Body Ownership (BO). Despite long-standing philosophical and scientific interest in the relationship between the body and brain, the neural systems involved in SoA and BO and especially their interactions, are not yet understood. In this preregistered study using the Moving Rubber Hand Illusion inside an MR-scanner, we aimed to uncover the relationship between BO & SoA in the human brain. Importantly, by using both visuomotor and visuotactile stimulations and measuring online trial-by-trial fluctuations in the illusion magnitude, we were able to disentangle brain systems related to objective sensory stimulation and subjective judgments of the bodily-self. Our results indicate that at both the behavioral and neural levels, BO and SoA are strongly interrelated. Multisensory regions in the occipital and fronto-parietal regions encoded convergence of sensory stimulation conditions. However, the subjective judgments of bodily-self were also related to BOLD fluctuations in regions not activated by the sensory conditions such as the insular cortex and precuneus. Our results highlight the convergence of multisensory processing in specific neural systems for both BO and SoA with partially dissociable regions for subjective judgments in regions of the default mode network.

## Introduction

A fundamental aspect of human phenomenology is the sensation of being an embodied agent in the world (Blanke & Metzinger, 2009; De Vignemont, 2018; Limanowski & Blankenburg, 2013; Seth & Tsakiris, 2018). Our experience in and of the world is constructed through our bodily sensations and actions, which as William James pointed out, “is always there” (James, 1890). The basic embodied sense of selfhood, termed Bodily Self-Consciousness, comprises two primary aspects. Body Ownership (BO), the experience of identifying with a body (Blanke, 2012; Ehrsson et al., 2004; Salomon et al., 2013; Tsakiris, 2010), and the Sense of Agency (SoA), the feeling of control over one’s actions (Gallagher, 2007; Haggard, 2017; Krugwasser et al., 2019; Moore & Fletcher, 2012). Under normal conditions, both BO and SoA are phenomenologically transparent and considered pre-reflexive (De Vignemont, 2018; Gallagher, 2000; Haggard, 2017), forming the implicit basis for our sense of self (Blanke & Metzinger, 2009; Gallagher, 2000; Salomon, 2017). Predictive processing accounts (Clark, 2013; Friston, 2018; Limanowski & Friston, 2018) have been proposed to explain the formation of transparent self-models. These posit that the brain functions as a hierarchical inference machine endeavoring to predict sensory states based on prior experience (Allen & Friston, 2016; Clark, 2013; Friston, 2018; Rao & Ballard, 1999). Such accounts thus propose that the self is formed through predictive models of the integration of interoceptive (Park et al., 2016; Park & Blanke, 2019; Salomon, Ronchi, et al., 2016; Seth, 2013), exteroceptive (Blanke et al., 2015; Botvinick & Cohen, 1998; Gentile et al., 2013; Limanowski, 2022) and volitional (Chambon et al., 2014; Haggard, 2008; Hara et al., 2015; Stern et al., 2022; Tsakiris et al., 2010) signals, proposed to underlie both BO (Chancel et al., 2021; Ehrsson & Chancel, 2019; Samad et al., 2015) and SoA (Constant et al., 2022; Legaspi & Toyoizumi, 2019; Leptourgos & Corlett, 2020).

Experimental investigations have highlighted the role of multisensory integration in the formation of BO (e.g., Blanke et al., 2015; Gentile et al., 2013). For example, in the classical Rubber Hand Illusion (RHI) paradigm (Botvinick & Cohen, 1998), viewing touch on a rubber hand while receiving anatomically and temporally synchronous tactile stimulation on one’s unseen hand creates a sense of illusory ownership over the rubber hand (Aimola Davies et al., 2013; Costantini & Haggard, 2007; for review of methods see Riemer et al., 2019; Tsakiris, 2010). This illusory ownership over fake limbs or even full bodies is accompanied by changes in physiological measures such as galvanic skin response (GSR), responses to threats (Armel & Ramachandran, 2003; Ehrsson et al., 2007), changes in skin temperature (Moseley et al., 2008; but see Rohde et al., 2013; Salomon et al., 2013), proprioceptive drift (Aimola Davies et al., 2013; Costantini & Haggard, 2007) and motor evoked potentials (Della Gatta et al., 2016), indicating that the illusion impacts not only explicit subjective experience but also physiological processing. Thus, BO is assumed to arise from the unconscious integration of interoceptive and exteroceptive sensory signals (Blanke et al., 2015; Chancel et al., 2021; Park & Blanke, 2019; Salomon, 2017; Seth, 2013). At the neural level, experimental studies in neurological patients and illusory ownership paradigms in healthy participants have revealed the involvement of frontal brain regions such as ventral premotor cortex (PMv), occipito-temporal regions such as anterior insula (AI), extrastriate body area (EBA), posterior regions such as posterior parietal cortex (PPC), and intraparietal sulcus (IPS), as well as the Cerebellum (Grivaz et al., 2017; Petkova et al., 2011; Salvato et al., 2020; Seghezzi, Giannini, et al., 2019; Tsakiris, 2010).

The sense of agency is known to rely on multisensory integration but also utilizes internal volitional signals (Haggard, 2008). The “comparator model” suggests that actions are accompanied by forward models predicting their sensory outcomes (Blakemore et al., 2002; Wolpert et al., 1995; but see Carruthers, 2012). These predictions, are then compared to actual sensory signals and when they are congruent, SoA arises (Frith, 2012; Jeannerod, 2009). Numerous studies have shown that when a conflict is introduced between the predicted outcomes of an action and the actual outcomes, SoA is reduced (e.g., Krugwasser et al., 2019; Sato, 2009; Stern et al., 2020). For example, when the spatial (Kannape et al., 2010; Nielsen, 1963; Salomon et al., 2022; Yomogida et al., 2010), anatomical (Krugwasser et al., 2019; Salomon, Fernandez, et al., 2016), or temporal (Farrer et al., 2008; Koreki et al., 2015; Limanowski et al., 2017; Stern et al., 2020; Wen et al., 2015) consequences of an action are deviated, this causes a reduction of SoA ratings. Importantly, we make a distinction between SoA experiments employing non-embodied paradigms, in which the sensory outcome of an action is learned during the experiment (Aarts et al., 2005; Sato, 2009; Sidarus et al., 2013; Wen et al., 2015; Yomogida et al., 2010) and embodied SoA studies, in which the expectations are based on lifelong prior experience with sensorimotor contingencies (Kalckert & Ehrsson, 2012; Krugwasser et al., 2022; Ma & Hommel, 2015; Salomon et al., 2022; Stern et al., 2020; Tsakiris et al., 2010). The neural systems underlying embodied SoA have been challenging to study due to the intricacies arising from executing, tracking, and altering bodily actions within neuroimaging environments. Previous studies using both embodied and non-embodied SoA paradigms point to the involvement of frontal motor regions such as the PMv, supplementary motor area (SMA and pre-SMA), as well as the Insula and occipito-temporal regions such as the PPC and the cerebellum (Farrer et al., 2007; Limanowski et al., 2017; Salomon et al., 2009; Tsakiris et al., 2010; for reviews see Haggard, 2017; Seghezzi, Giannini, et al., 2019; Sperduti et al., 2011).

The relation between BO and SoA has been hotly debated. Several accounts suggest that they are dissociable phenomena (Gallagher, 2000; Kalckert & Ehrsson, 2012; Seghezzi, Giannini, et al., 2019). Neurological deficits, for example, may impact one aspect or the other suggesting a dissociation between BO and SoA. While ‘somatoparaphrenia’ patients show a change in BO experiencing a loss of ownership over their hand (Feinberg & Venneri, 2014; Vallar & Ronchi, 2009). In ‘Anarchic Hand Syndrome’, the patients show loss of SoA while retaining BO (Biran & Chatterjee, 2004). Historically, experimental studies of BO and SoA were typically separated, limiting our understanding of their interdependencies. BO has typically been studied using RHI-type paradigms in which the illusory ownership of a fake hand or body is induced through synchronous visuotactile stimulation (e.g., Botvinick & Cohen, 1998; Costantini & Haggard, 2007; Riemer et al., 2019). SoA on the other hand, has most often been studied by inducing sensorimotor conflicts in which the visual or auditory outcomes deviates from the predicted outcome (e.g., Farrer et al., 2008; Krugwasser et al., 2019; Ma & Hommel, 2015; Wen et al., 2015). Recently, a new paradigm named the Moving Rubber Hand Illusion (MRHI) (Kalckert & Ehrsson, 2012, 2014), allows to module both visuomotor (VM) and visuotactile (VT) correspondences and is thus suited to test both BO and SoA within the same participant and session. Despite several decades of research, the neural correlates of BO and SoA and especially their interdependencies are not well understood. Critically, previous RHI studies associated the brain activity during illusion eliciting conditions (synchronous and congruent VT or VM conditions) with a subjective report of a similar condition before or after the imaging session (e.g., Ehrsson et al., 2005; Gentile et al., 2013; Ionta et al., 2014; Tsakiris et al., 2010) As such, online, trial-by-trial variance in the subjective experiences of BO and SoA are overlooked and BO & SoA were equated with the multisensory conditions (e.g., synchronous or asynchronous stimulation) that give rise to the illusions.

In this preregistered study (https://aspredicted.org/KRJ_WPI) we used an adaptation of the MRHI during fMRI scanning. We manipulated VT and VM correspondences in the temporal and anatomical domains to alter the experience of BO and SoA. Importantly, we measured both BO and SoA on each trial allowing to (1) investigate the subjective and neural interplay between BO and SoA; (2) compare neural processing of different sensorimotor conflicts, and (3) to disentangle the brain activity related to multisensory stimulation from that related to the subjective experience of BO and SoA. Specifically, if BO and SoA are independent phenomena, they should be dissociated at both the behavioral and neural levels. Conversely, if they are correlated in behavior and show overlapping neural systems, this would suggest that they are interrelated. Additionally, we hypothesized that brain regions tracking subjective BO and SoA judgments outside the networks activated by the multisensory stimulation are related to the phenomenological experience of BO and SoA.

## Materials and methods

### Participants

Thirty healthy right-handed subjects participated in the experiment (9 female), between 19 and 37 years of age (M = 26.8, SD = 4.4). All participants gave written informed consent before the experiment, and the study was approved by the ethics committee of Be’er Ya’akov’s mental health center and corresponded to the human subject’s guidelines of the declaration of Helsinki. The number of participants was based on our preregistered power analysis.

Preregistered exclusion criteria included previous or current neurological or psychiatric disorder, medication use, tactile or motor deficits, non-normal or non-corrected to normal vision, and contraindications that prohibited magnetic resonance imaging (MRI) scanning. We recruited only participants who are ≥168cm tall to ensure that their hand is accessible for tactile stimulation in the constrained space of the scanner environment. Post-scan exclusion criteria included subjects with significant cerebral abnormalities, enlarged ventricles, cysts, and severe cerebral asymmetry, as well as runs with excessive head movements during the scan (>3.5mm). Prior to the analysis of any contrast of interest, excessively noisy runs were also excluded from analysis (see preregistration for full details). Due to these exclusion criteria, 3 subjects were excluded from the functional analysis along with another 15 runs.

### Experimental setup

To familiarize the participants with the paradigm and the MRI environment, the participants underwent a brief pre-scan single run inside a mock MRI scanner, including all VT and VM conditions. Only in the pre-scan participants were instructed to press a control pad with their left hand when they first started to feel “as if the rubber hand was their own” (experience of BO) to indicate the estimated timing for the onset of the illusion in the scanner. After having made the keypress response, participants were instructed to continue maintaining their gaze at the fixation cross dyed on the rubber hand. This data was not analyzed in the scope of the present study. In both pre-scan and scanning phases, participants rested comfortably in a supine position on the bed, with their right arm extended and placed on a support in a relaxed position. All participants were wearing headphones to reduce noise and to receive auditory cues. Two fMRI compatible mirrors were placed above the participants’ faces to ensure they could see the apparatus with a natural view. We used an adaptation of the MRHI setup previously described in (Kalckert & Ehrsson, 2012, 2014). Participants’ right hand was placed into a tilted box (20 cm × 15 cm × 10 cm) located on the participant’s right side of their waist (see Fig. 1A). A realistic life-sized and gender-matched hand model was placed on top of the box and covered with a latex glove. Subjects wore identical latex gloves on their right hands, and their forearm was covered with a soft black cloth to ensure visual continuity of the rubber hand with the participant’s arm. Since VT stimulation has been suggested to affect results through the experimenter’s seen movements (Limanowski et al., 2014), the stimulation of brushstrokes was done with long wooden brushes that allowed the experimenter to remain outside the subject’s field of view. In VM runs, we used technical Lego parts and a regular rhythm of 1 Hz to simplify the control for the number of movements. To help the participants and the experimenter apply the same number of movements or brushstrokes in the VM or VT runs, both listened to an auditory metronome at 1 Hz over earphones. In all VT and VM conditions, the experimenter monitored the number of movements to verify if missed stimulations or movements occurred. To keep the subjects’ gaze on the rubber hand, the rubber hand was marked with a fixation cross on the back of the palm above both fingers. A similar cross was marked on the glove of the subject’s real hand.

**Figure 1.**
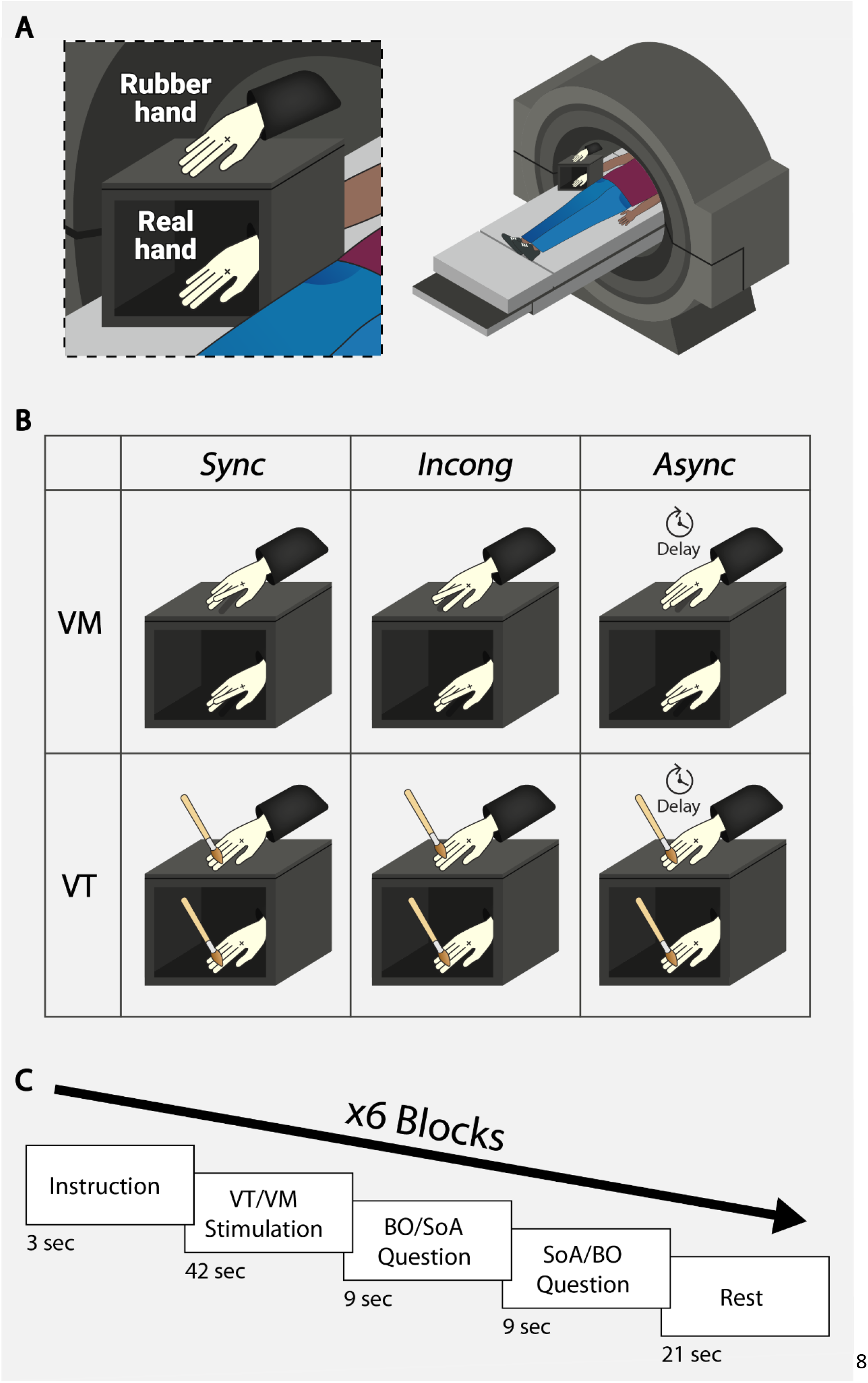
Experimental setup and design. **(A)** Experimental setup of the moving rubber hand illusion paradigm inside the MR-scanner. **(B)** Experimental design illustration of VM and VT stimulations each divided for *Sync, Incong*, and *Async* conditions. **(C)** Timeline of a run including VM or VT stimulation and a subjective rating for BO and SoA.

### Experimental design

fMRI runs were divided into separate VT and VM runs. Each run had 6 blocks with three types of conditions: Synchronous Congruent (*Sync*), Synchronous Incongruent (*Incong*), and Asynchronous Congruent (*Async*). In both VT and VM runs, brush strokes or an active movement of the participant’s index finger accompanied the same stimulation or movement in the rubber hand index finger synchronously (*Sync*), the rubber hand middle finger (*Incong*), or the index finger but with a 500ms delay between hands (*Async*) (see Fig. 1B). In VT blocks, participants were instructed to relax their hand while the experimenter brushed both hands with small brushes and manipulated the position or timing of brush strokes between them. In VM blocks, participants were instructed to raise their index finger at 1 Hz after each auditory cue of the metronome. The index finger and middle finger of the rubber hand could be mechanically connected to the participants’’ index finger, allowing them to be yoked to the participants’ movements (*Sync* and *Incong*) or to the experimenter’s unseen hand (*Async*) using a thin long stick. Both runs began with 30 seconds of rest followed by 6 pseudorandomized blocks, containing two of each three types of conditions (*Sync, Incong, Async*). Each block began with 3 seconds of auditory instructions (‘Passive’ or ‘Active’) reminding the participant whether the upcoming trial would involve VT or VM conditions (see Fig. 1C). Following the instructions, each block contained stimulation of brushstrokes or active movements, which lasted 42 seconds. Immediately after the end of each condition, the subjects were instructed to rate their experience of BO and SoA (by hearing the words ‘Ownership’ and ‘Agency’) towards the rubber hand. Each rating had a limited time of 9 s in which the participants responded using an fMRI-compatible response pad containing 4 buttons (ranging from completely agree to completely disagree). The ratings’ order was pseudorandomized between subjects. Following each block, participants had a rest period of 21 seconds in which they were instructed to close their eyes.

### fMRI data acquisition

The scan sessions were performed in a Siemens Prisma 3T MRI scanner, equipped with an 18-channel head coil. Whole-brain functional images were acquired using an echo planar imaging (EPI) sequence with AC-PC alignment (TR=3000ms, TE=30ms, matrix size: 490 × 490, flip angle: 85° in-plane resolution: 3mm × 3mm × 3mm slice, no gap). Interleaved slices were acquired in an ascending direction. In total, 173 volumes were acquired throughout each run (1038 volumes in total), such that a single fMRI run covers the duration of each of the single behavioral runs. T1-weighted anatomical scans were acquired for each participant (192 slices; voxel resolution: 1 × 1 × 1mm, no gap; TR: 2300ms; TE: 2.32ms, inversion time: 900ms, flip angle of 8°, matrix size: 256 × 256).

### Behavioral data analysis

Participants’ behavioral ratings did not pass the Shapiro-Wilk normality test and were therefore analyzed by the nonparametric Friedman test to assess the alterations of BO and SoA experiences across conditions (*Sync, Incong, Async*). To examine if a condition induced an illusion of BO or SoA over the rubber hand, we conducted a Wilcoxon signed rank test by comparing the participants’ ratings to 0 (i.e., neutral experience) and corrected for multiple comparisons using false discovery rate (FDR). The differences between the BO and SoA ratings during VM and VT were analyzed using Wilcoxon rank-sum test. We performed the nonparametric Bayesian versions of these tests to evaluate null findings using JASP (JASP Team, 2019). To evaluate the association of BO and SoA ratings, we correlated the participants’ illusion scores by subtracting each *Sync* rating from the average baseline rating in the *Async* condition for both BO and SoA.

### fMRI data preprocessing and analysis

Imaging data were preprocessed and analyzed using BrainVoyager 21.4 software (Brain Innovation, Maastricht, The Netherlands). All functional imaging data underwent the same series of preprocessing before all successive analyses. The functional volumes were motion corrected to the first volume of each series, corrected for slice-timing errors by cubic spline interpolation. Low-frequency signal drifts in the images were removed by a high-pass filter (2 sines/cosines). The functional volumes were co-registered to the high-resolution structural image and were spatially smoothed with a 6 mm FWHM Gaussian kernel. Each participant’s functional images were segmented into gray matter, white matter, and cerebrospinal fluid (CSF) partitions and were normalized to the MNI standard space. A general linear model (GLM) was fitted for each participant with regressors modeling the three types of conditions (*Sync, Incong, Async*). All maps were corrected for multiple comparisons using a Monte Carlo Cluster-Level Statistical Threshold Estimator (Forman et al., 1995) with the Cluster Threshold estimator plugin (Brainvoyager, Brain Innovation) at alpha=0.005. Cluster-size voxel threshold was set according to the estimation for each map: VM *Sync>Async* - 54 voxels, VM *Sync>Incong* - 51 voxels, VT *Sync>Async* - 28 voxels and VT *Sync>Incong* - 29 voxels.

Maps were then converted to cluster VOIs using BrainVoyager’s “Convert Map Clusters to VOI(s)” function with the same cluster-size threshold used to correct the map for multiple comparisons. Cluster coordinates were extracted using both cluster “Peak” table and “Center of Gravity” table functions in BrainVoyager’s VOI Analysis option menu and were used as inputs to determine clusters’ anatomical names using AAL (Automated Anatomical Labeling) and suggested BA (Brodmann area) proposed by “Label4MRI” R package (https://github.com/yunshiuan/label4MRI), (see Supplementary Tables S3-S10).

### Regions of interest localization

All regions of interest (ROIs) were pre-defined in our preregistration. In an additional scanning session at the end of all experimental runs, we employed a functional localizer for identifying standard motor and sensory regions that were not analyzed in the present paper. Our pre-registered ROIs were defined by the overlap of *Sync>Async* VT activation clusters with the regions defined in previous fMRI studies RHI (see Fig. 4A and Supplementary Table S3). Note that we then investigated VM block activity in these ROIs, thus ensuring complete independence between ROI definition and analysis of signals within the ROIs.

### Correlations of ROIs with subjective ratings

BO and SoA scores were averaged for each condition (*Sync, Incong, Async*) in each run (VM and VT), and correlated with the mean beta value within our pre-defined ROIs. These correlations were tested in a Wilcoxon signed rank test at each region by comparing the participants’ correlations to zero. The direction of the test was defined for positive correlations in activated regions and negative correlations in deactivated regions (R TPJ).

### Whole brain parametric GLMs of Subjective BO and SoA ratings

The trial-by-trial subjective ratings for BO and SoA were used as subject-specific parametric regressors for both VM and VT runs. To ensure a balanced model, we included only subjects who had three successful runs in VM or VT and that their ratings differed at least once within each run, which resulted with four parametric maps: VM SoA (n=22), VT SoA (n=19), VM BO (n=23), and VT BO (n=20). All maps were corrected for multiple comparisons at alpha<0.05, cluster-size voxel threshold was set according to estimation for each map: VM SoA - 252 voxels, VM BO - 200 voxels, VT SoA - 174 voxels and VT BO - 111 voxels.

### Psychophysiological Interaction (PPI)

We used a PPI analysis (Friston et al., 1997) to examine which brain regions change their connectivity with the Insula during the illusion (*Sync*), compared with non-illusory conditions (*Async+Incong*). First, we independently defined the left and right Insula as a seed region based on the Harvard-Oxford atlas (https://identifiers.org/neurovault.collection:262). For each seed region, a standard psychophysiological (PPI) analysis was carried out for each subject using the PPI plugin for Brainvoyager (V1.30) on the VM trials.

In detail, the contrast of interest was defined as *Sync>*(*Incong+Async*), and weights were given accordingly, while non-relevant conditions such as instructions and fixation were set to zero. After defining the contrast of interest, the time course (BOLD signal) of the seed region (left and right Insula, atlas defined) of each subject was extracted, Z-transformed, and then convolved with the hemodynamic response function. Next, TR was multiplied by TR and with the task time course (based on the protocol associated with the data) to create the PPI predictor (the interaction term). In addition to the PPI predictor, three additional predictors were created: a psychological regressor, based on the task protocol, a physiological regressor, which is the seed region time course and a complementary regressor. Next, for each seed region, a second-level analysis was done using multi-subject GLM analysis with the predictors described above. Maps show clusters of voxels in which there were specific changes in the connectivity with the Insula, suggesting an exchange of information. All PPI maps were FDR corrected at q<0.05.

### Deviations from preregistered analysis plan

We have deviated from the preregistered analysis plan in the following: Representational Similarity Analysis of the data (preregistered analysis # 6) and functional connectivity analysis for rest data (preregistered analysis # 9) are beyond the scope of the current manuscript. The PPI analysis was finally conducted on the VM vs. VT instead of the BO vs. SoA. Additionally, the partition of data based on the onset of the illusion (preregistered analysis # 3) could not be conducted as the onset of the illusion in the VM condition was so rapid that it often did not allow comparisons of pre-post fMRI data.

## Results

### Subjective experience of Body Ownership and Sense of Agency

Illusory BO over the rubber hand was modulated by the experimental conditions for both VM (Friedman test; χ2(2) = 44.9, *p*<0.0001) and VT (Friedman test; χ2(2) = 41.3, *p*<0.0001) stimulations. Similarly, SoA ratings were modulated by the conditions in both VM (Friedman test; χ2(2) = 44.4, *p*<0.0001) and VT (Friedman test; χ2(2) = 45.8, *p*<0.0001). As expected, the *Sync* condition induced an illusion of ownership over the rubber hand in both VM (Wilcoxon signed rank test; M = 0.57, SD = 0.46, V = 387, *p*<0.0001; FDR corrected) and VT (Wilcoxon signed rank test; M = 0.57, SD = 0.45, V = 438, *p*<0.0001; FDR corrected) stimulations (see Fig. 2A and B). Similarly, the *Sync* condition induced an experience of SoA in VM (Wilcoxon signed rank test; M = 0.77, SD = 0.34, V = 461, *p*<0.0001; FDR corrected) and VT (Wilcoxon signed rank test; M = 0.31, SD = 0.54, V = 324, *p*<0.01; FDR corrected) stimulations. Importantly, only the *Sync* condition induced an illusion of BO and SoA significantly above zero. All other conditions revealed substantially more evidence for the null hypothesis (all BF10 < 0.092) apart from the SoA ratings in the VM *Incong* condition showing an intermediate effect (BF10 = 0.74) (see Supplementary Table S1). Critically, and as expected, VM elicited a stronger main effect of SoA ratings than the passive VT (Wilcoxon rank-sum test; W = 378, p<0.001; FDR corrected). These results are driven by the higher SoA ratings in the *Sync* condition (Wilcoxon rank-sum test; W = 310, p<0.0001; FDR corrected) and the *Incong* condition (Wilcoxon rank-sum test; W = 395.5, p<0.0001; FDR corrected) in VM compared to VT stimulations. Importantly, no significant difference was found for the main effect of BO (p>0.05, BF10 = 0.092) or the SoA ratings in the *Incong* and *Async* conditions (see Supplementary Table S2). Next, we tested correlations between BO and SoA ratings and found highly significant correlations between BO and SoA ratings in both VM (r=0.41, p<0.05, *Spearman*) and VT (r=0.65, p<0.001, *Spearman*) stimulations (see Fig. 2C and D).

**Figure 2.**
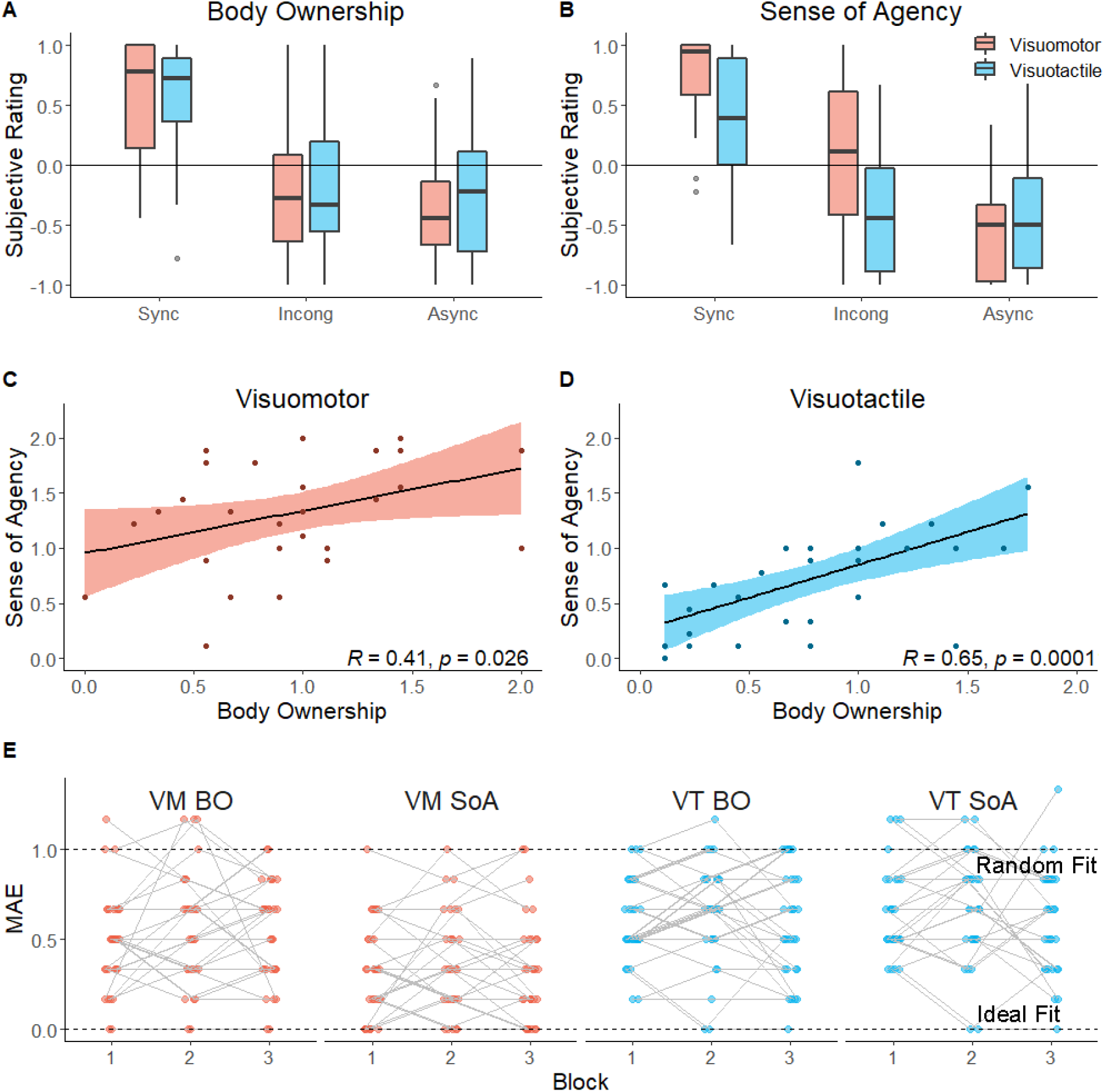
Behavioral results. **(A)** BO and **(B)** SoA subjective ratings during VM and VT (blue) stimulations. **(C)** Correlation between BO and SoA ratings during VM stimulation **(D)** and VT stimulation. **(E)** MAE of BO and SoA ratings. The graph illustrates the deviation of subjective rating from an ideal model (lower dashed line) and a random model (upper dashed line).

### Fluctuations in the experience of Body Ownership and Sense of Agency

We examined the stability of the experience of the illusion across time and subjects by calculating the Mean Absolute Error (MAE) from an “ideal model” represented by the highest subjective rating for *Sync* and the minimum rating for *Async* (see Fig. 2E) and a “random model” denoting equal chances for high or low subjective ratings for these conditions. Across subjects, BO experience ratings differed significantly from the ideal model for both VM (Wilcoxon signed rank test; M = 0.52, SD = 0.23, V = 406, p<0.0001; FDR corrected) and VT (Wilcoxon signed rank test; M = 0.60, SD = 0.23, V = 465, p<0.0001; FDR corrected). Similarly, SoA experience ratings in VM (Wilcoxon signed rank test; M = 0.33, SD = 0.24, V = 406, p<0.0001; FDR corrected) and VT (Wilcoxon signed rank test; M = 0.63, SD = 0.23, V = 465, p<0.0001; FDR corrected) differed considerably from this ideal model, confirming considerable inter-subject variability in susceptibility to the MRHI illusion. To investigate the variations of subjective ratings across different instances of the illusion within each subject, we calculated the mean absolute differences of these errors between blocks. The results indicated that the within subjective ratings differed across blocks for the BO experience ratings in VM (Wilcoxon signed rank test; M = 0.21, SD = 0.18, V = 325, p<0.0001; FDR corrected) and VT (Wilcoxon signed rank test; M = 0.18, SD = 0.11, V = 351, p<0.0001; FDR corrected) and as well as for SoA experience ratings in VM (Wilcoxon signed rank test; M = 0.15, SD = 0.13, V = 325, p<0.0001; FDR corrected) and VT (Wilcoxon signed rank test; M = 0.19, SD = 0.15, V = 300, p<0.0001; FDR corrected). This result indicates that significant intra-subject variability in the experience of the illusion is also present in the data.

### Neural correlates of the Bodily Self

#### Temporal conflict processing

As per our preregistered analysis, we first examined if similar brain regions were involved in the processing of illusory states produced by VM and VT stimulations. We produced whole-brain maps of *Sync*>*Async*, as this is the condition associated with the modulations of both BO and SoA for both VM and VT stimulations.

For VM stimulation (Fig. 3A), the contrast map of the *Sync>Async* revealed increased BOLD responses in posterior occipital regions extending dorsally to parietal and frontal regions. Left hemisphere sensorimotor regions were activated, in line with movements of the right hand. In the right hemisphere, activations were also present along the superior bank of the STS, and notably, deactivations (i.e., higher BOLD response in the *Async*) were present in the inferior parietal and insular regions. Medial regions, including the precuneus and SMA, were also more active in the *Sync* condition.

**Figure 3.**
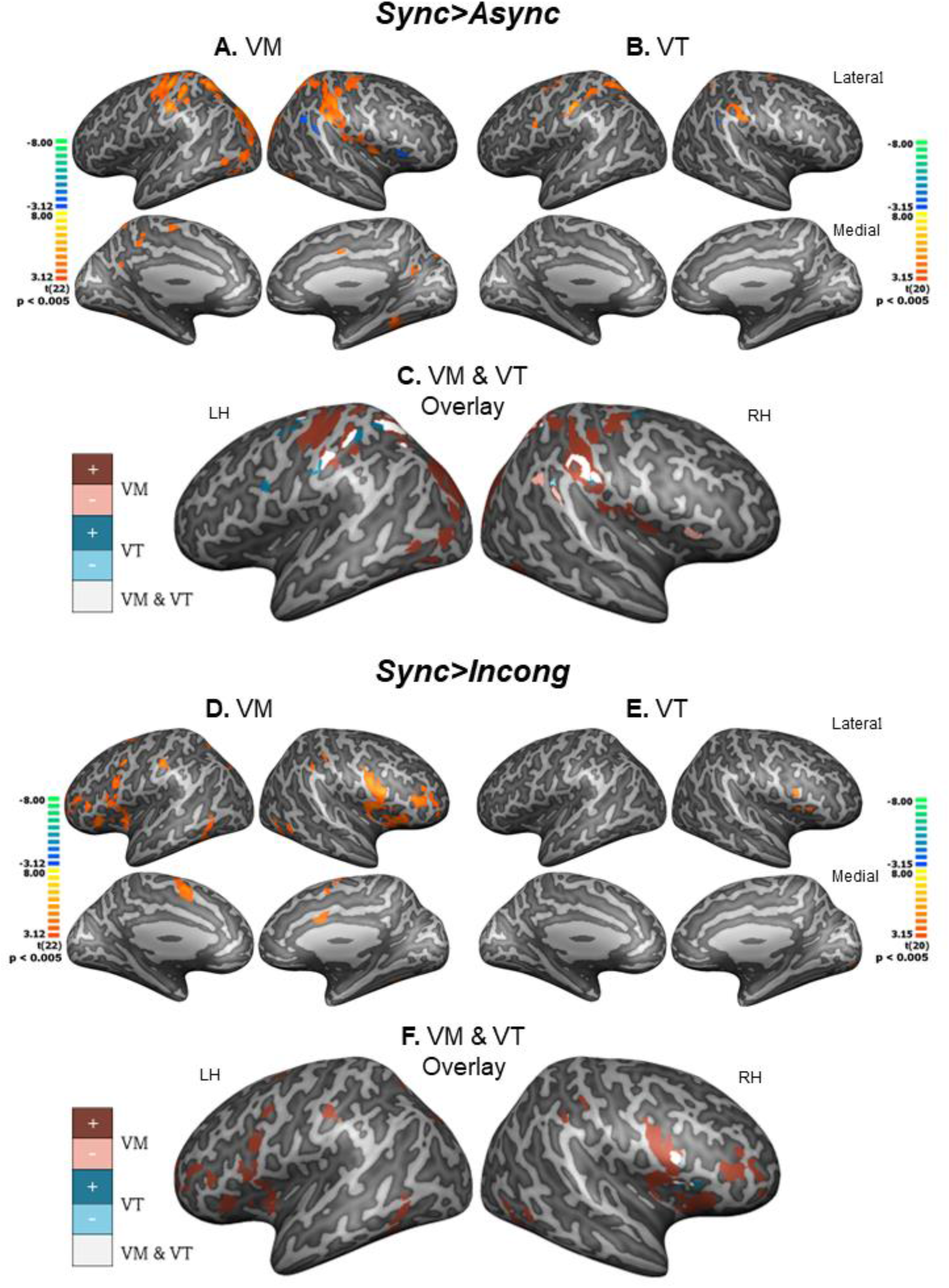
Whole brain analysis of bodily multisensory conflicts. **(A)** VM *Sync>Async* contrast map. **(B)** VT *Sync>Async* contrast map. **(C)** VM&VT *Sync>Async* contrast maps overlaid and displayed in solid colors (VM: dark red and pink, VT: blue and light blue), dark colored clusters were more activated during *Sync* condition while lighter colored clusters were more activated during *Async*, overlapping clusters are displayed in white. **(D)** VM *Sync>Incong* contrast map. **(E)** VT *Sync>Incong* contrast map. **(F)** VM&VT *Sync>Incong* contrasts overlaid, dark colored clusters were more activated during *Sync* condition while lighter colored clusters were more activated during *Incong*, overlapping clusters are displayed in white. All contrast maps are displayed in neurological orientation, thresholded at p<0.005, and corrected for multiple comparisons.

During VT stimulation, the *Sync>Async* contrast map at the same threshold elicited less significant activity (Fig. 3B). As can be seen the *Sync* condition elicited a higher BOLD response than the *Async* condition in parietal regions of the intraparietal sulcus, dorsal posterior parietal cortex, and post central sulcus, as well as frontal premotor regions. A small patch of deactivation was found in the right inferior parietal cortex. The overlap of these two maps (Fig. 3C) shows that regions of the bilateral Posterior Parietal cortex (PPC) and bilateral postcentral were more active during the *Sync* stimulation across both VM and VT stimulation conditions.

#### Anatomical conflict processing

We also wished to investigate anatomical sensorimotor conflicts in which the temporal correlation is maintained, but the movement or tactile stimulation are on the middle finger (i.e., *Incong* condition). This condition also reduced BO and SoA ratings, but it is unclear if this conflict is associated with similar brain regions as those processing temporal conflicts as in the *Async* condition.

During VM stimulation (Fig. 3D), the activity map of the *Sync>Incong* contrast revealed increased BOLD responses across lateral inferior frontal regions, including the insular cortex and extending dorsally to premotor regions. Bilaterally, a region in the posterior parietal cortex and in the post central sulcus. Left hemisphere precentral and postcentral sulcus regions were activated, with more robust BOLD responses during *Sync* compared to *Incong* anatomical feedback. Medial regions, including the ACC and SMA were also more active in the *Sync* condition, and lateral occipital regions. During VT stimulation, the *Sync>Incong* contrast elicited less significant activity (Fig. 3E). As can be seen, the *Sync* condition elicited higher BOLD response than the *Incong* condition in the right insular cortex and frontal operculum. As can be seen in figure 3F, this region showed an overlap between the VM and VT stimulations (see Supplementary Fig. S1 for further comparisons).

#### Preregistered Region of Interest analysis

As per our preregistration, we wished to investigate the overlap of sensorimotor conflict processing (*Sync>Async*) in several regions of interest (see Fig. 4A). As can be seen in figure 4B across the eight pre-registered ROIs, five (L IPS, L LOC, L PPC, L Somatosensory and L PMv) showed significant effects similar to those found in the VT (i.e., stronger activity in the *Sync* condition than the *Async* condition) and one (R TPJ) showed similar increased activity in the *Async* conditions. The SMA and L insular regions did not show significant differences. Thus, across most regions of interest temporal conflicts induced a differential BOLD response regardless of the stimulation method.

**Figure 4.**
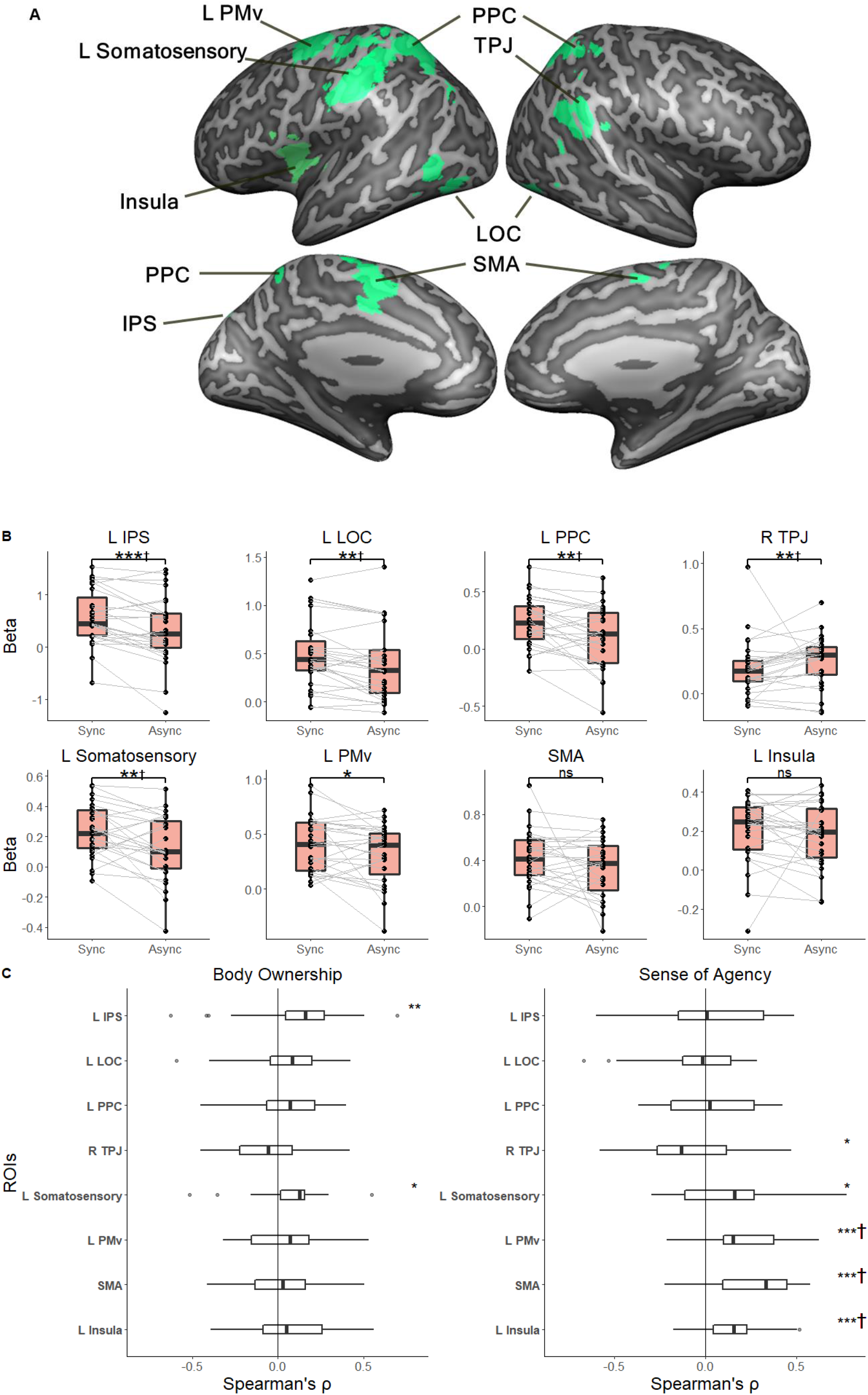
BO and SoA correlation with the brain activity in pre-defined ROIs. **(A)** Eight preregistered ROIs were defined using VT stimulation blocks. **(B)** Mean beta values of each ROI were extracted for VM *Sync* and *Async* conditions and compared using paired Wilcoxon signed-rank test. **(C)** Correlation analysis between the BO and SoA ratings in the scanner with the brain activity scores across pre-defined ROIs are presented as boxplot distributions of the subject’s correlation within each ROI. Asterisks indicate significance levels: \**p*<0.05; \*\**p*<0.01; and \*\*\**p*<0.001. † indicate FDR corrected results.

In order to assess if our regions of interest reflect subjective judgments of SoA and BO we examined the correlation of BO and SoA scores with the mean beta value within our pre-defined ROIs (Fig. 4A). The correlation of reported BO was positively and significantly correlated with the beta values within the L IPS (V = 285, p<0.01) and the L Somatosensory (V = 263, p<0.05). In addition, SoA ratings were significantly and positively correlated with the beta values within the L Somatosensory (V = 270, p<0.05), the L PMv (V = 358, p<0.0001, FDR corrected), the SMA (V = 353, p<0.0001, FDR corrected), the L Insula (V = 342, p<0.0001, FDR corrected), and significantly negatively correlated within the R TPJ (V = 112, p<0.05).

### Whole brain correlates of subjective experience of BO and SoA

We examined neural systems relating to the subjective experiences of BO and SoA through parametric GLMs using trial-by-trial SoA and BO ratings. As can be seen in Figures 5A-C, the ratings of SoA were associated in both the VT and VM stimulations with sensorimotor regions predominantly in the left hemisphere. Medial regions extending from frontal SMA to posterior medial and lateral parietal regions (bilateral PPC) were positively associated with SoA ratings, as were lateral occipital regions (Bilateral LOC and L IPS). Interestingly, the insular cortex bilaterally was also activated during high SoA trials. Additionally, and as expected, SoA activation in the VM task included additional regions, including medial frontal regions, anterior cingulate, and the precuneus.

**Figure 5.**
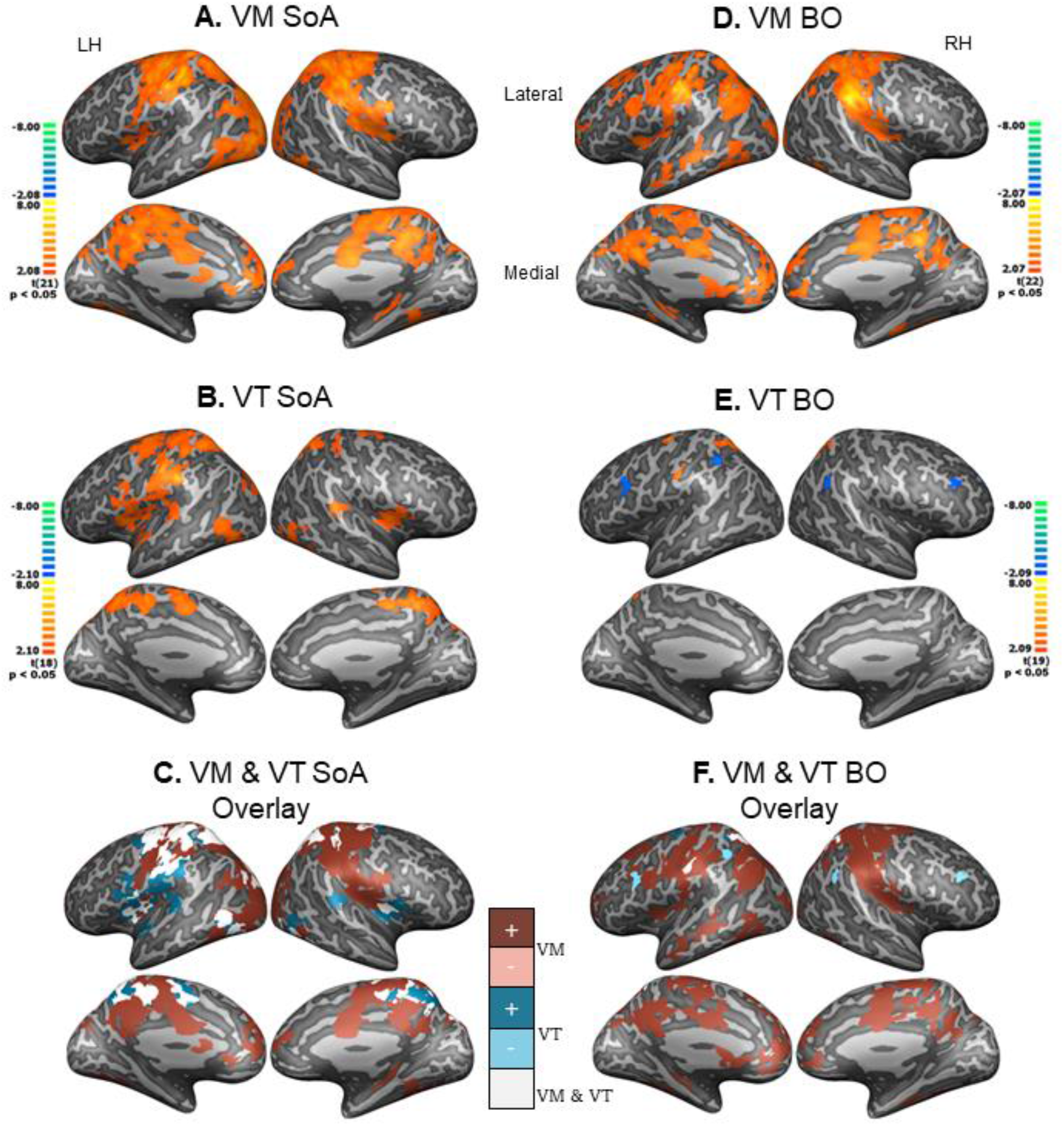
Neural correlates of subjective Sense of Agency and Body Ownership. **(A)** VM SoA parametric map. **(B)** VT SoA parametric map. **(C)** VM&VT SoA overlaid and displayed in solid colors (VM: dark red and pink, VT: blue and light blue), dark colored clusters were related to the SoA while lighter colored clusters are inversely related to it, overlapping clusters are displayed in white. **(D)** VM BO parametric map. **(E)** VT BO parametric map. **(F)** VM & VT BO overlaid dark colored clusters were related to BO while lighter colored clusters were inversely related to it, overlapping clusters are displayed in white. All maps were thresholded at p<0.05 and corrected for multiple comparisons.

Parametric mapping of BO experience (Fig 5D-F) revealed that during VM stimulation, more expansive regions of the cortex were associated with subjective ratings than in the VT stimulation using the same statistical threshold. In the VM blocks, BO was associated with higher activity in sensorimotor and parietal regions extending along the temporal sulcus. Interestingly, BO activated medial regions including the precuneus, ACC and SMA and extending to midline prefrontal regions (BA 8,10,32).

During VT blocks, the parametric mapping of BO was much more localized and included a region of the left premotor cortex, left somatosensory cortex, and the dorsal posterior parietal regions. The latter two regions overlapped with BO regions from the VM blocks. Two additional small patches negatively associated with BO rating during VT blocks were found in the TPJ and frontal regions (BA12 and BA9 respectively).

#### Cerebellar activations

Although not part of our preregistered ROIs, *Sync> Async* in both VT and VM strongly activated regions of the cerebellum, and these were also correlated with subjective ratings of BO and SoA (see Supplementary Fig. S3).

### Insula Psychophysical Interaction analysis

Given the insular’s cortex involvement in BO and SoA and its suggested role in predictive coding, we used the Insula as a seed region for a PPI analysis. We compared the *Sync* condition in VM stimulation with the non-illusion conditions *Async* and *Incong*. Comparison of connectivity between these conditions revealed that during the *Sync* condition, the Insula was more correlated with the left sensorimotor, ACC, lateral and medial prefrontal regions, as well as lateral occipital regions (Fig. 6A-B and supplementary Tables S9-10). These findings were similar for both right and left Insula, though the left Insula was correlated with a larger number of voxels. Thus, during the illusion eliciting condition, the insular cortex was more tightly correlated with widespread regions primarily in the frontal and occipital cortices.

**Figure 6.**
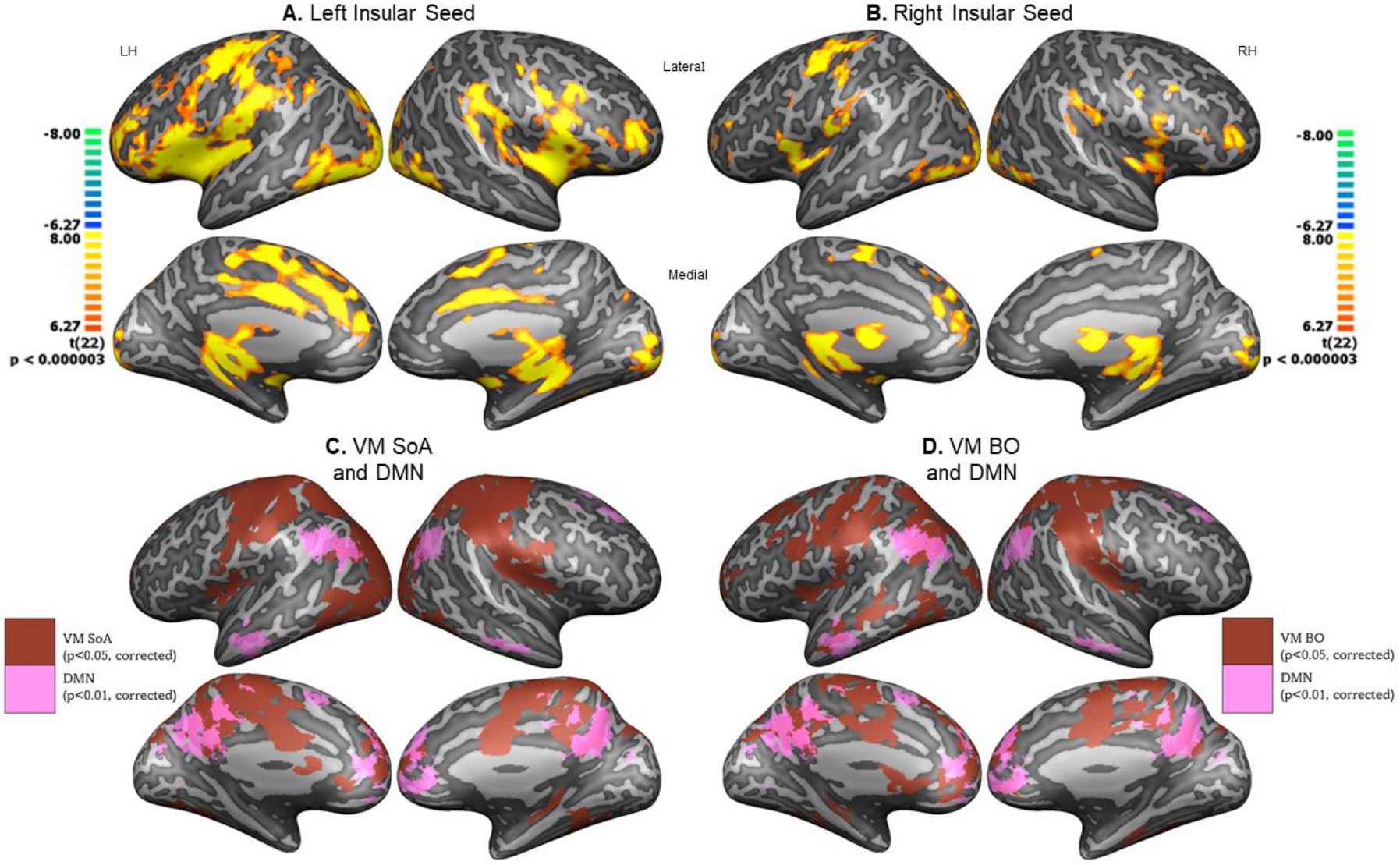
Psychophysiological Interaction analysis from insular seed & Default Mode Network overlap. Task-based effective connectivity from insular regions during VM stimulation for illusion condition (*Sync>Incong+Async*), using left **(A)** and right **(B)** Insula. Group maps were FDR corrected at q<0.05. Note: Illusory condition was associated with stronger connectivity between Insula and left sensorimotor, ACC and LOC. DMN overlap with subjective experience **(C)** VM SoA (same as Fig. 5A) and the DMN as defined by Neurosynth.org (“DMN-association test”, FDR corrected q=0.05). **(D)** VM BO (same as Fig. 5D) and the DMN. Note: subjective judgements of SoA and BO considerably overlap with medial components of the DMN (Precuneus and vmPFC).

## Discussion

Our preregistered study aimed to investigate the relationship between the neural systems underlying the subjective sensations of SoA and BO and their relation to the Visuotactile and Visuomotor stimulations which elicit them. The results revealed several interesting findings. Behaviorally, in line with previous work, we found that the subjective ratings of BO and SoA were significantly correlated, suggesting an interdependent relationship between the two. Furthermore, while temporally synchronous and anatomically congruent VT and VM stimulations elicited illusory ownership and agency, trial-by-trial measurements showed that participants had considerable variability in their subjective experiences. Analysis of fMRI data indicated (1) extensive overlap in the brain systems related to stimulation inducing illusory bodily experiences. (2) Differential activations were elicited by temporal and anatomical conflicts. (3) The brain systems underlying trial-by-trial subjective effects of BO and SoA showed overlaps in sensorimotor, insular, and parietal regions. However, subjective sensations of the bodily self arising from volitional actions were associated with activity in regions of the DMN.

### Behavioral and neural convergence of bodily multisensory conflict processing

The theoretical separation of BO and SoA has allowed increased conceptual clarity to the study of the bodily self. Experimental, neuropsychological, and imaging studies have shown that these may be dissociated both behaviorally and in their underlying neural systems (Gallagher, 2000; Seghezzi, Giannini, et al., 2019; Tsakiris et al., 2010). However, recent experimental paradigms testing BO and SoA in unison (Kalckert & Ehrsson, 2014) have revealed correlations between the two, suggesting that these are not independent under normal conditions. Conceptually, the ability to control the action of a limb would provide strong evidence that it is “my own” and should induce a feeling of ownership over it (Salomon, Fernandez, et al., 2016; Tsakiris et al., 2010). Our behavioral results show that across both VM and VT stimulation methods, BO and SoA were strongly correlated. Indeed, even when no actions were performed, passive VT stimulation caused participants to feel agency over the rubber hand. At the neural level, six of eight preregistered ROIs, selected and localized from VT illusion condition showed a similar significant difference in the VM illusion condition (Fig. 3). Thus, most brain regions encoding bodily multisensory conflicts across the occipito-parietal cortices do so for both passive VT conflicts as well as VM conflicts. Whole brain mapping of illusion eliciting conditions revealed similar results with considerable overlap and adjacent activity for *Sync* across VM and VT stimulations in PPC, IPS, and TPJ regions (Fig. 3C). Thus, *Sync* stimulation of a rubber hand from both tactile and motor origins activates multisensory parietal regions as well as occipital and frontal sensorimotor regions. This finding extends previous work showing recruitment of frontal-parietal multisensory regions during the classical VT rubber hand illusion (Ehrsson et al., 2005; Gentile et al., 2013; Limanowski et al., 2014; Tsakiris et al., 2010) indicating their involvement in active volitional action as in the VM MRHI condition.

### Divergent neural processing for temporal and anatomical conflicts

To test the specificity of multisensory conflict processing, we included an anatomical conflict in which the temporal synchrony was maintained, but the incorrect finger moved. Interestingly, in VM, the anatomical conflict (*Sync*>*Incong*) was associated primarily with reduced activity in frontal regions extending from the insular cortex through the frontal operculum as well as R TPJ. During VT stimulation, the same contrast elicited less significant activations converging in the insular cortex (Fig. 3D-F). A direct comparison of the conflict conditions (*Async-Incong*) revealed similar results with reduced activation in frontal regions during the anatomical conflict and reduced activations in the parietal regions for temporal conflicts. Critically, in the VM condition there was an overlap of temporal and anatomical conflict processing in the LOC, Somatosensory and insular regions (Supplementary Fig. S1). This results extends previous reports of regions of LOC responding to one’s actions (Astafiev et al., 2004), and VT multisensory congruence (David et al., 2007; Limanowski et al., 2014), bodily illusions (Apps et al., 2015; Gentile et al., 2013; Limanowski et al., 2014) to processing of errors arising from both temporal and anatomical conflicts. Despite the convergence of conflict processing in these three regions, our findings highlight a frontal-parietal divergence for different bodily prediction errors. It is possible that computations of low-level temporal deviations are processed in classical multimodal integration regions across the dorsal parietal regions known to be related to VM processing (Blanke, 2012; Grivaz et al., 2017; Seghezzi, Giannini, et al., 2019). However, anatomical deviations which are “categorical” errors and uncommon in real life, may be computed in more frontal regions relating to high level prediction errors (Alexander & Brown, 2018; Corlett et al., 2022).

In the PPI analysis, examination of the connectivity of the Insula for the *Sync* conditions compared with the conflict conditions, showed increased coupling between the Insula, ACC, sensory motor regions and LOC suggesting that temporal and anatomical conflicts decrease exchange of information and the correlation between them.

### Dissociating sensory stimulation and subjective experience

Most brain imaging examinations during the RHI have typically used offline measurements of subjective experience as correlates for activity during imaging. This approach has not allowed to tease apart activity related to objective sensory stimulations from activity related to the subjective experiences of BO and SoA. Here we measured trial-by-trial fluctuations of subjective experiences while maintaining constant sensory stimulation conditions. Indeed, we found highly significant variability in participants’ experiences of BO and SoA for a given sensory stimulation (Fig. 2E). Thus, parametric mapping of these online modulations of phenomenological experience allowed us to decouple objective sensory stimulations from the subjective experiences of SoA and BO.

Preregistered ROIs showed significant correlations with subjective reports. As per our preregistered prediction, frontal regions related to sensorimotor processing (L PMv, SMA, and L Somatosensory cortex) and the L insular cortex were positively correlated with SoA. These frontal regions have been previously related to the processing of SoA (David et al., 2008; Haggard, 2017; Salomon et al., 2009; Sperduti et al., 2011; Yomogida et al., 2010). Specifically, the SMA and PMv have been suggested to have a central role in comparing action intentions and sensory outcomes to allow adaptive monitoring of motor control (Seghezzi, Zirone, et al., 2019; Sperduti et al., 2011). Thus, it has been suggested that these regions (as well as the cerebellum) are part of the neural circuitry of the “comparator model” continuously matching the action’s sensory predictions with afferent signals (Blakemore et al., 2001; Haggard, 2017; Sperduti et al., 2011). Indeed, these regions and especially the PMv and somatosensory regions were also activated in the illusory stimulation condition, suggesting that they are associated with both sensorimotor prediction error as well as subjective aspects of SoA judgements.

Whole brain analysis of the trial-by-trial subjective experience of SoA showed that in both VT and VM frontal-parietal regions, including the Insula, PMV, Sensorimotor regions extending to the PPC and LOC were associated with higher SoA ratings (Fig. 5C). While many of these regions were also activated by the illusory stimulation condition (*Sync>Async* see Supplementary Fig. S2 for overlap), the left Insula, ACC and precuneus regions were activated only during subjective judgements of SoA. While the insular cortex has been previously related to SoA processing (Seghezzi, Giannini, et al., 2019; Seghezzi, Zirone, et al., 2019; Sperduti et al., 2011; Tsakiris et al., 2010), bodily self (Grivaz et al., 2017; Seghezzi, Giannini, et al., 2019; Tsakiris et al., 2007) and interoception (Critchley & Harrison, 2013; Quadt et al., 2018; Salomon, Ronchi, et al., 2016), our results reveal that this is driven by subjective judgments of SoA rather than multisensory conflict processing.

Importantly, midline structures such as the ACC have been found to be involved in self-related processing (Qin et al., 2010, 2020; Wittmann et al., 2021). Experimental and Meta Analytical work has shown the ACC has been related to processes such as self-face and self-referential processing (Hu et al., 2016; Ulmer-Yaniv et al., 2021), self-other interactions (Lockwood et al., 2018; Shimon-Raz et al., 2021; Wittmann et al., 2021) and self-monitoring (Iannaccone et al., 2015; Shenhav et al., 2016). The ACC and insular cortex are also considered the core elements of the “Salience” network suggested to guide attention between internal and external directed cognition (Uddin, 2015). Our data shows that subjective judgments of SoA recruit the ACC and Insula, suggesting that the subjective decision process may involve a shift of attention from external stimulation to internal mentation typically associated with Default Mode network regions.

When participants made volitional movements (VM), subjective judgements of SoA and BO were associated with activity in midline regions of the DMN including VMPFC and the precuneus (Fig. 6C-D). These regions of the DMN are associated with self-related processing (Andrews-Hanna et al., 2014; Davey et al., 2016; Peer et al., 2015; Salomon et al., 2014) across various manipulations and paradigms (for meta-analysis see Qin & Northoff, 2011). Critically, in our paradigm, subjective SoA and BO judgements during VM conditions activated medial prefrontal and precuneus regions of the DMN (for overlap with DMN see Fig. 6). This finding, links sensorimotor and multisensory regions typically associated with the bodily self (Blanke, 2012; Haggard, 2017; Seghezzi, Giannini, et al., 2019) to DMN regions relating to more conceptual levels of self-processing (Peer et al., 2015; Qin et al., 2020; Salomon et al., 2014). Thus, we propose that while BO and SoA components of the bodily self are driven through multisensory integration of visual, bodily, motor, and interoceptive signals in frontoparietal networks, the phenomenological aspect recruits regions of the DMN. We note that this is specific to the VM condition, suggesting that the involvement of efferent self-generated actions is essential for linking bodily and conceptual levels of the self. This idea is supported by a plethora of evidence linking volitional actions to enhanced self-representation in perception (Salomon, Szpiro-Grinberg, et al., 2011; Wen et al., 2018; Wen & Imamizu, 2022), memory (Pacheco Estefan et al., 2021; Rotem-Turchinski et al., 2019), and consciousness (Suzuki et al., 2019). Moreover, numerous studies have shown that deficits in SoA in psychiatric populations with aberrant self-representations such as in the Schizophrenia spectrum (Hauser et al., 2011; Krugwasser et al., 2022; Salomon et al., 2022; Synofzik et al., 2010) who also show deviant DMN connectivity (Camchong et al., 2009; Salomon, Bleich-Cohen, et al., 2011; Dong et al., 2018).

In summary, our study has allowed a novel investigation of the neural mechanisms underlying SoA and BO and their relation to the multisensory stimulations giving rise to them. By tracking trial-by-trial fluctuations in subjective experience, we were able to show converging and disparate systems involved in the multisensory processes and the subjective experience revealing a central role for the insular cortex and midline structures of the DMN in the experience of the Bodily Self.

## Acknowledgements

The authors thank the MRI team at the Weizmann institute for science and Prof. Rafi Malach for their help with this study. This study was supported by an Israeli Science Foundation personal grant (ISF 1169/17) to R.S.

