## Supplemental Information for "Disentangling the Neural Correlates of Agency, Ownership and Multisensory Processing"

### Table S1 - Illusion effect

| Stimulation | Experience | Condition | n | mean | sd | V | p | FDR | $BF_{10}$ |
|---|---|---|---|---|---|---|---|---|---|
| Visuomotor | Body Ownership | Sync | 30 | 0.578 | 0.464 | 387 | 1.2e-05 | † | 1965.993 |
| Visuomotor | Body Ownership | Incong | 30 | -0.230 | 0.524 | 96 | 0.99 | | 0.065 |
| Visuomotor | Body Ownership | Async | 30 | -0.378 | 0.451 | 51.5 | 1 | | 0.084 |
| Visuomotor | Sense of Agency | Sync | 30 | 0.770 | 0.349 | 461.5 | 8.9e-07 | † | 23101.875 |
| Visuomotor | Sense of Agency | Incong | 30 | 0.130 | 0.615 | 269 | 0.13 | | 0.747 |
| Visuomotor | Sense of Agency | Async | 30 | -0.552 | 0.380 | 8 | 1 | | 0.114 |
| Visuotactile | Body Ownership | Sync | 30 | 0.578 | 0.452 | 438 | 1.2e-05 | † | 51.069 |
| Visuotactile | Body Ownership | Incong | 30 | -0.207 | 0.506 | 124.5 | 0.99 | | 0.089 |
| Visuotactile | Body Ownership | Async | 30 | -0.207 | 0.523 | 112 | 0.98 | | 0.081 |
| Visuotactile | Sense of Agency | Sync | 30 | 0.315 | 0.540 | 324.5 | 0.0029 | † | 6365.578 |
| Visuotactile | Sense of Agency | Incong | 30 | -0.459 | 0.486 | 28 | 1 | | 0.071 |
| Visuotactile | Sense of Agency | Async | 30 | -0.407 | 0.482 | 52.5 | 1 | | 0.070 |

**Table S1 - Illusion effect -** The illusion effect of BO and SoA was examined by comparing the participants' ratings to 0 (i.e., neutral experience) in a Wilcoxon signed rank test, corrected for multiple comparisons using FDR. Significant effects are marked in green. Significant FDR corrected results are marked with †. Bayes factors were calculated with Bayesian Wilcoxon signed rank test.

Table S2 - Differences between Visuomotor and Visuotactile ratings

| Experience | Condition | W | p | FDR | $BF_{10}$ |
|---|---|---|---|---|---|
| Body Ownership | Sync | 105.5 | 0.5 | | 0.207 |
| Body Ownership | Incong | 134 | 0.68 | | 0.164 |
| Body Ownership | Async | 100 | 0.98 | | 0.067 |
| Sense of Agency | Sync | 310 | 3.7e-05 | † | 1819.408 |
| Sense of Agency | Incong | 395.5 | 6.2e-05 | † | 1064.694 |
| Sense of Agency | Async | 122 | 0.92 | | 0.073 |

**Table S2 - Differences between Visuomotor and Visuotactile ratings -** The differences between the BO and SoA ratings during VM and VT were analyzed using Wilcoxon rank-sum test, corrected for multiple comparisons using FDR. To evaluate all null findings, we performed a Bayesian Wilcoxon rank-sum test. Significant effects are marked in green. Significant FDR corrected results are marked with †.

Table S3 - Preregistered ROIs

| | Peak Activation | | | | | | | Center of Gravity | | | | | | | | |
|---|---|---|---|---|---|---|---|---|---|---|---|---|---|---|---|---|
| | | | | | | | | Mean | | | STD | | | | | |
| ROI Name | X | Y | Z | t | p | Anatomical Labeling | Brodmann Area | X | Y | Z | X | Y | Z | #Voxels | Anatomical Labeling | Brodmann Area |
| L IPS | -15 | -68 | 61 | 2.627966 | 0.016565 | Precuneus_L | Left-BA7 | -20.09 | -78.24 | 52.47 | 7.93 | 4.94 | 7.22 | 1967 | Parietal_Sup_L | Left-BA7 |
| L LOC | -51 | -70 | -5 | 3.342205 | 0.003424 | Occipital_Inf_L | Left-BA19 | -48.64 | -71.04 | -2.24 | 5.32 | 4.03 | 3.01 | 2195 | Occipital_Inf_L | Left-BA19 |
| L PPC | -30 | -52 | 55 | 4.134723 | 0.000563 | Parietal_Inf_L | Left-BA7 | -27.26 | -51.31 | 57.44 | 7.66 | 7.51 | 6.29 | 8566 | Parietal_Sup_L | Left-BA7 |
| R TPJ | 49 | -46 | 39 | -3.435242 | 0.002774 | Parietal_Inf_R | Right-BA40 | 52.44 | -40.80 | 30.54 | 6.48 | 3.53 | 5.15 | 4008 | SupraMarginal_R | Right-BA40 |
| L Somatosensory | -60 | -19 | 34 | 3.346418 | 0.003391 | Postcentral_L | Left-PrimSensory (1) | -46.63 | -27.71 | 43.09 | 9.67 | 6.12 | 8.76 | 7509 | Parietal_Inf_L | Left-BA40 |
| L PMv | -30 | -16 | 67 | 5.582672 | 0.000022 | Precentral_L | Left-BA6 | -28.30 | -12.11 | 61.52 | 7.63 | 5.65 | 7.30 | 9332 | Precentral_L | Left-BA6 |
| SMA | -6 | -7 | 55 | 2.288298 | 0.033752 | Supp_Motor_Area_L | Left-BA6 | -3.69 | -6.19 | 51.46 | 4.11 | 4.78 | 6.90 | 3271 | Cingulum_Mid_L | Left-BA24 |
| L Insula | -39 | -1 | 10 | 2.642563 | 0.016055 | Insula_L | Left-Insula (13) | -40.94 | -4.08 | 8.01 | 3.81 | 4.99 | 5.02 | 2953 | Insula_L | Left-Insula (13) |

**Table S3 – Preregistered ROIs** - MNI coordinates of ROIs (by “Peak Activity” and “Center of Gravity”) and their Automated Anatomical Labeling and Brodmann Area (BA) output by “Label4MRI” R package.

Table S4 - Visuotactile significant clusters

| | Peak Activation | | | | | | | Center of Gravity | | | | | | | | |
|---|---|---|---|---|---|---|---|---|---|---|---|---|---|---|---|---|
| | | | | | | | | Mean | | | STD | | | | | |
| Contrast Map | X | Y | Z | t | p | Anatomical Labeling | Brodmann Area | X | Y | Z | X | Y | Z | #Voxels | Anatomical Labeling | Brodmann Area |
| VT Sync > Async | 60 | -16 | 31 | 4.764917 | 0.000118 | Postcentral_R | Right-PrimSensory (1) | 54.36 | -19.45 | 35.82 | 4.19 | 3.34 | 5.00 | 1465 | Postcentral_R | Right-PrimSensory (1) |
| VT Sync > Async | 33 | -46 | 61 | 4.478062 | 0.000230 | Parietal_Sup_R | Right-BA7 | 28.78 | -50.02 | 58.19 | 5.76 | 6.59 | 3.59 | 951 | Parietal_Sup_R | Right-BA7 |
| VT Sync > Async | 15 | -58 | -47 | 4.952794 | 0.000077 | Cerebelum_8_R | Right-Fusiform (37) | 14.83 | -62.92 | -46.64 | 4.30 | 6.07 | 2.63 | 1458 | Cerebelum_8_R | Right-Fusiform (37) |
| VT Sync > Async | -27 | -52 | 55 | 5.747705 | 0.000013 | Parietal_Inf_L | Left-BA7 | -28.32 | -47.31 | 53.88 | 6.95 | 8.43 | 4.60 | 3512 | Parietal_Inf_L | Left-BA7 |
| VT Sync > Async | -30 | -16 | 61 | 4.426879 | 0.000259 | Precentral_L | Left-BA6 | -32.66 | -13.28 | 61.81 | 3.43 | 3.39 | 6.90 | 1474 | Precentral_L | Left-BA6 |
| VT Sync > Async | -54 | -22 | 40 | 6.711431 | 0.000002 | Parietal_Inf_L | Left-BA40 | -54.00 | -22.97 | 38.33 | 4.56 | 3.02 | 3.17 | 1405 | Parietal_Inf_L | Left-BA40 |
| VT Sync > Incong | 54 | 11 | 16 | 4.176417 | 0.000466 | Frontal_Inf_Oper_R | Right-BA44 | 50.36 | 15.25 | 14.26 | 3.70 | 5.29 | 5.42 | 2080 | Frontal_Inf_Oper_R | Right-BA44 |
| VT Sync > Incong | 27 | 35 | -2 | 5.007642 | 0.000068 | Frontal_Inf_Orb_R | Right-BA47 | 27.37 | 32.76 | -2.86 | 3.36 | 5.54 | 3.22 | 1440 | Frontal_Inf_Orb_R | Right-BA47 |
| VT Async > incong | 60 | -43 | 34 | 4.513497 | 0.000212 | SupraMarginal_R | Right-BA40 | 57.97 | -40.00 | 32.81 | 2.70 | 4.24 | 5.06 | 862 | SupraMarginal_R | Right-BA40 |
| VT All conditions | -54 | -22 | 40 | 6.711431 | 0.000002 | Parietal_Inf_L | Left-BA40 | -2.75 | -38.01 | 12.15 | 33.98 | 41.40 | 31.71 | 691049 | Cingulum_Post_L | Left-BA30 |
| VT All conditions | 21 | -16 | -20 | 2.479093 | 0.022201 | ParaHippocampal_R | Right-Parahip (36) | 38.45 | -17.50 | -13.43 | 12.16 | 17.58 | 13.68 | 23825 | Hippocampus_R | Right-Hippocampus (54) |
| VT All conditions | 30 | 65 | -35 | 1.893066 | 0.072911 | Frontal_Mid_Orb_R | Right-BA10 | 36.38 | 59.61 | -39.18 | 6.34 | 5.68 | 4.23 | 5498 | Frontal_Mid_Orb_R | Right-BA10 |
| VT All conditions | -33 | -20 | 57 | 3.000582 | 0.007067 | Precentral_L | Left-BA6 | 7.50 | -19.41 | 49.60 | 21.64 | 11.19 | 11.84 | 21633 | Supp_Motor_Area_R | Right-BA6 |
| VT All conditions | -6 | -16 | 28 | -2.306053 | 0.031944 | Cingulum_Mid_L | Left-BA23 | 0.54 | -29.07 | 27.69 | 5.02 | 8.56 | 4.36 | 6566 | Cingulum_Mid_R | Right-BA23 |
| VT All conditions | -57 | -1 | -5 | 2.498898 | 0.021284 | Temporal_Sup_L | Left-BA22 | -35.14 | -25.57 | -5.88 | 13.45 | 19.91 | 13.50 | 17223 | Hippocampus_L | Left-Caudate (48) |
| VT All conditions | -29 | 59 | -26 | 2.407439 | 0.025841 | Frontal_Mid_Orb_L | Left-BA10 | -34.44 | 58.50 | -38.16 | 7.10 | 6.27 | 4.75 | 7720 | Frontal_Inf_Orb_L | Left-BA10 |

**Table S4 - Visuotactile significant clusters -** MNI coordinates of clusters extracted from VT stimulation contrast maps (p<0.005, corrected) and their suggested Automated Anatomical Label and Brodmann Area (BA) output by "Label4MRI" R package.

Table S5 - Visuomotor significant clusters

| | Peak Activation | | | | | | | Center of Gravity | | | | | | | | |
|---|---|---|---|---|---|---|---|---|---|---|---|---|---|---|---|---|
| | | | | | | | | Mean | | | STD | | | | | |
| Contrast Map | X | Y | Z | t | p | Anatomical Labeling | Brodmann Area | X | Y | Z | X | Y | Z | #Voxels | Anatomical Labeling | Brodmann Area |
| VM Sync > Async | 27 | -76 | 40 | 5.644993 | 0.000011 | Occipital_Sup_R | Right-BA7 | 36.87 | -46.91 | 41.06 | 13.45 | 28.92 | 16.71 | 36493 | Parietal_Inf_R | Right-BA7 |
| VM Sync > Async | 45 | -61 | -8 | 4.654527 | 0.000122 | Temporal_Inf_R | Right-Fusiform (37) | 47.79 | -62.98 | -9.72 | 6.96 | 5.98 | 4.09 | 3238 | Temporal_Inf_R | Right-Fusiform (37) |
| VM Sync > Async | 21 | -46 | -23 | 4.447120 | 0.000202 | Cerebelum_4_5_R | Right-Fusiform (37) | 26.45 | -44.13 | -17.67 | 5.50 | 5.91 | 5.21 | 3637 | Cerebelum_4_5_R | Right-Fusiform (37) |
| VM Sync > Async | 0 | -10 | 46 | 4.145200 | 0.000423 | Cingulum_Mid_L | Right-BA24 | -0.52 | -9.11 | 46.64 | 3.48 | 7.24 | 5.88 | 2709 | Cingulum_Mid_L | Left-BA24 |
| VM Sync > Async | -21 | -85 | 34 | 5.842810 | 0.000007 | Occipital_Sup_L | Left-BA19 | -24.07 | -82.33 | 27.70 | 7.71 | 6.39 | 10.57 | 14335 | Occipital_Sup_L | Left-BA19 |
| VM Sync > Async | -36 | -31 | 55 | 6.597615 | 0.000001 | Postcentral_L | Left-PrimMotor (4) | -32.99 | -33.07 | 54.54 | 10.50 | 13.49 | 8.49 | 18272 | Postcentral_L | Left-PrimSensory (1) |
| VM Sync > Async | -30 | -52 | -17 | 4.160712 | 0.000407 | Fusiform_L | Left-Fusiform (37) | -32.19 | -51.47 | -17.91 | 7.25 | 3.27 | 4.08 | 1829 | Fusiform_L | Left-Fusiform (37) |
| VM Sync > Async | -48 | -73 | 4 | 4.302758 | 0.000288 | Occipital_Mid_L | Left-BA19 | -42.14 | -72.65 | -2.74 | 5.52 | 3.85 | 4.33 | 2047 | Occipital_Inf_L | Left-BA19 |
| VM Sync > Incong | 51 | 14 | 19 | 5.457166 | 0.000018 | Frontal_Inf_Oper_R | Right-BA44 | 41.42 | 20.04 | 4.93 | 12.80 | 14.52 | 12.19 | 25557 | Frontal_Inf_Tri_R | Right-BA45 |
| VM Sync > Incong | 51 | -67 | 1 | 4.597924 | 0.000140 | Temporal_Mid_R | Right-Fusiform (37) | 50.72 | -60.86 | -3.99 | 3.81 | 7.58 | 5.67 | 2859 | Temporal_Inf_R | Right-Fusiform (37) |
| VM Sync > Incong | -6 | 2 | 49 | 4.838927 | 0.000078 | Supp_Motor_Area_L | Left-BA6 | -1.99 | -0.49 | 55.81 | 7.11 | 5.19 | 9.06 | 4854 | Supp_Motor_Area_L | Left-BA6 |
| VM Sync > Incong | -21 | 59 | 16 | 4.601221 | 0.000139 | Frontal_Sup_L | Left-BA10 | -28.04 | 53.79 | 10.01 | 6.74 | 4.15 | 5.45 | 2662 | Frontal_Mid_L | Left-BA10 |
| VM Sync > Incong | -39 | -4 | 43 | 5.302387 | 0.000025 | Precentral_L | Left-BA6 | -45.88 | 5.34 | 12.75 | 9.37 | 5.94 | 15.58 | 8680 | Rolandic_Oper_L | Left-BA44 |
| VM Sync > Incong | -39 | 26 | 7 | 6.196434 | 0.000003 | Frontal_Inf_Tri_L | Left-BA45 | -36.07 | 29.87 | 8.10 | 5.67 | 6.02 | 6.53 | 4314 | Frontal_Inf_Tri_L | Left-BA45 |
| VM Sync > Incong | -42 | -52 | -11 | 4.880857 | 0.000070 | Fusiform_L | Left-Fusiform (37) | -44.56 | -57.24 | -7.32 | 5.26 | 5.70 | 5.29 | 2089 | Temporal_Inf_L | Left-Fusiform (37) |

**Table S5 - Visuomotor significant clusters -** MNI coordinates of clusters extracted from VM stimulation contrast maps (p<0.005, corrected) and their suggested Automated Anatomical Label and Brodmann Area (BA) output by "Label4MRI" R package.

Table S6 - Visuomotor significant clusters

| | Peak Activation | | | | | | | Center of Gravity | | | | | | | | |
|---|---|---|---|---|---|---|---|---|---|---|---|---|---|---|---|---|
| | | | | | | | | Mean | | | STD | | | | | |
| Contrast Map | X | Y | Z | t | p | Anatomical Labeling | Brodmann Area | X | Y | Z | X | Y | Z | #Voxels | Anatomical Labeling | Brodmann Area |
| VM Async > Incong | 66 | -40 | 25 | 5.673763 | 0.000010 | SupraMarginal_R | Right-BA22 | 59.43 | -39.00 | 25.87 | 6.00 | 4.72 | 4.69 | 4507 | SupraMarginal_R | Right-BA40 |
| VM Async > Incong | 60 | -49 | 4 | 5.707229 | 0.000010 | Temporal_Mid_R | Right-Fusiform (37) | 57.33 | -52.82 | 6.35 | 4.04 | 6.46 | 3.02 | 1833 | Temporal_Mid_R | Right-Fusiform (37) |
| VM Async > Incong | 42 | 26 | 1 | 6.076862 | 0.000004 | Frontal_Inf_Tri_R | Right-BA45 | 37.59 | 18.84 | 9.68 | 13.05 | 14.92 | 16.43 | 27947 | Frontal_Inf_Oper_R | Right-BA44 |
| VM Async > Incong | 54 | -43 | 52 | 4.232515 | 0.000342 | Parietal_Inf_R | Right-BA40 | 50.78 | -48.46 | 46.18 | 4.83 | 4.62 | 4.45 | 2058 | Parietal_Inf_R | Right-BA40 |
| VM Async > Incong | 30 | -82 | 34 | -4.645901 | 0.000125 | Occipital_Mid_R | Right-BA19 | 26.82 | -80.45 | 38.86 | 4.75 | 3.70 | 5.72 | 2358 | Occipital_Sup_R | Right-BA7 |
| VM Async > Incong | 21 | -58 | 55 | -5.005958 | 0.000052 | Parietal_Sup_R | Right-BA7 | 19.14 | -57.95 | 59.92 | 3.06 | 3.15 | 3.68 | 1510 | Parietal_Sup_R | Right-BA7 |
| VM Async > Incong | 9 | -82 | -20 | 4.427812 | 0.000212 | Cerebelum_Crus1_R | Right-VisualAssoc (18) | 12.27 | -79.76 | -26.40 | 4.26 | 3.05 | 6.97 | 2001 | Cerebelum_Crus1_R | Right-VisualAssoc (18) |
| VM Async > Incong | 3 | 14 | 55 | 6.461407 | 0.000002 | Supp_Motor_Area_R | Right-BA6 | 3.25 | 8.13 | 60.69 | 7.33 | 10.15 | 7.26 | 8578 | Supp_Motor_Area_R | Right-BA6 |
| VM Async > Incong | -15 | -82 | 37 | -5.082895 | 0.000043 | Cuneus_L | Left-BA19 | -17.55 | -82.04 | 36.69 | 4.78 | 3.73 | 4.67 | 2656 | Occipital_Sup_L | Left-BA19 |
| VM Async > Incong | -12 | -76 | -29 | 5.404827 | 0.000020 | Cerebelum_Crus1_L | Left-VisualAssoc (18) | -17.53 | -78.58 | -27.29 | 8.56 | 5.55 | 4.39 | 3891 | Cerebelum_Crus1_L | Left-VisualAssoc (18) |
| VM Async > Incong | -36 | 23 | 10 | 5.993311 | 0.000005 | Frontal_Inf_Tri_L | Left-BA45 | -36.80 | 19.59 | 7.03 | 8.69 | 9.84 | 8.35 | 11201 | Insula_L | Left-BA45 |
| VM Async > Incong | -36 | -31 | 55 | -6.744370 | 0.000001 | Postcentral_L | Left-PrimMotor (4) | -32.30 | -30.34 | 58.84 | 6.34 | 9.37 | 6.90 | 11471 | Postcentral_L | Left-PrimMotor (4) |
| VM Async > Incong | -27 | -64 | -47 | 4.882991 | 0.000070 | Cerebelum_8_L | Left-Fusiform (37) | -28.14 | -72.11 | -44.28 | 2.64 | 6.12 | 4.42 | 2150 | Cerebelum_Crus2_L | Left-BA19 |
| VM All conditions | 54 | -13 | -14 | -6.845551 | 0.000001 | Temporal_Mid_R | Right-BA22 | 55.59 | 5.70 | -11.52 | 12.44 | 21.61 | 17.88 | 26529 | Temporal_Pole_Sup_R | Right-BA38 |
| VM All conditions | -27 | -19 | -2 | 15.920792 | 0.000000 | Putamen_L | Left-Putamen (49) | -2.75 | -32.10 | 11.27 | 32.19 | 42.84 | 31.64 | 914181 | Cingulum_Post_L | Left-Thalamus (50) |
| VM All conditions | 42 | 53 | -41 | -11.385799 | 0.000000 | Frontal_Inf_Orb_R | Right-BA10 | 36.22 | 59.53 | -39.23 | 6.44 | 5.50 | 4.40 | 5528 | Frontal_Mid_Orb_R | Right-BA10 |
| VM All conditions | 18 | -37 | 64 | -7.137179 | 0.000000 | Postcentral_R | Right-PrimSensory (1) | 5.58 | -31.58 | 59.85 | 12.03 | 9.35 | 6.20 | 12329 | Paracentral_Lobule_R | Right-PrimMotor (4) |
| VM All conditions | 0 | 14 | -5 | -4.679196 | 0.000115 | Olfactory_L | Left-BA25 | 3.51 | 22.95 | -10.10 | 4.46 | 6.80 | 4.01 | 3323 | Frontal_Med_Orb_R | Right-BA25 |
| VM All conditions | -24 | 65 | -35 | -10.640593 | 0.000000 | Frontal_Sup_Orb_L | Left-BA11 | -34.49 | 58.36 | -38.14 | 7.08 | 6.46 | 4.91 | 7754 | Frontal_Inf_Orb_L | Left-BA10 |
| VM All conditions | -51 | -4 | -17 | -5.432136 | 0.000019 | Temporal_Mid_L | Left-BA21 | -51.77 | -6.65 | -14.59 | 4.26 | 8.21 | 5.84 | 2897 | Temporal_Mid_L | Left-BA22 |

**Table S6 - Visuomotor significant clusters -** MNI coordinates of clusters extracted from VM stimulation contrast maps (p<0.005, corrected) and their suggested Automated Anatomical Label and Brodmann Area (BA) output by "Label4MRI" R package.

Table S7 - Visuotactile whole brain correlates of the subjective experiences of Sense of Agency and Body Ownership

| | Peak Activation | | | | | | | Center of Gravity | | | | | | | | |
|---|---|---|---|---|---|---|---|---|---|---|---|---|---|---|---|---|
| | | | | | | | | Mean | | | STD | | | | | |
| Contrast Map | X | Y | Z | t | p | Anatomical Labeling | Brodmann Area | X | Y | Z | X | Y | Z | #Voxels | Anatomical Labeling | Brodmann Area |
| VT BO | 48 | -49 | 40 | -3.706413 | 0.001498 | Parietal_Inf_R | Right-BA39 | 46.16 | -42.16 | 30.80 | 8.97 | 7.30 | 11.65 | 4519 | SupraMarginal_R | Right-BA40 |
| VT BO | -15 | -46 | -20 | -3.927291 | 0.000905 | Cerebelum_4_5_L | Left-Fusiform (37) | -8.61 | -51.82 | -17.24 | 6.48 | 6.87 | 5.60 | 3416 | Cerebelum_4_5_L | Left-BA19 |
| VT BO | -30 | -52 | 55 | 4.134723 | 0.000563 | Parietal_Inf_L | Left-BA7 | -21.13 | -56.99 | 58.60 | 6.62 | 5.46 | 4.10 | 3819 | Parietal_Sup_L | Left-BA7 |
| VT BO | -30 | -16 | 67 | 5.582672 | 0.000022 | Precentral_L | Left-BA6 | -34.95 | -15.66 | 65.14 | 6.93 | 4.45 | 6.92 | 4573 | Precentral_L | Left-BA6 |
| VT SoA | 51 | -52 | -23 | 3.873907 | 0.001112 | Temporal_Inf_R | Right-Fusiform (37) | 52.51 | -50.03 | -1.68 | 9.03 | 16.65 | 14.07 | 15832 | Temporal_Mid_R | Right-Fusiform (37) |
| VT SoA | 24 | -64 | 64 | 5.460283 | 0.000035 | Parietal_Sup_R | Right-BA7 | -16.25 | -36.67 | 48.16 | 30.47 | 25.96 | 19.86 | 156250 | Cingulum_Mid_L | Left-SensoryAssoc (5) |
| VT SoA | 36 | 5 | 13 | 4.517231 | 0.000267 | Insula_R | Right-BA44 | 40.35 | 4.99 | 7.32 | 7.19 | 9.54 | 6.85 | 9047 | Insula_R | Right-BA44 |
| VT SoA | -24 | -61 | -20 | 4.060258 | 0.000734 | Cerebelum_6_L | Left-Fusiform (37) | -44.17 | -65.66 | -8.80 | 8.70 | 10.00 | 9.72 | 12151 | Occipital_Inf_L | Left-BA19 |

**Table S7 - Visuotactile whole brain correlates of the subjective experiences of Sense of Agency and Body Ownership -** MNI coordinates of clusters extracted from VT parametric maps (p<0.05, corrected) and their suggested Automated Anatomical Label and Brodmann Area (BA) output by "Label4MRI" R package.

Table S8 - Visuomotor whole brain correlates of the subjective experiences of Sense of Agency and Body Ownership

| | Peak Activation | | | | | | | Center of Gravity | | | | | | | | |
|---|---|---|---|---|---|---|---|---|---|---|---|---|---|---|---|---|
| | | | | | | | | Mean | | | STD | | | | | |
| Contrast Map | X | Y | Z | t | p | Anatomical Labeling | Brodmann Area | X | Y | Z | X | Y | Z | #Voxels | Anatomical Labeling | Brodmann Area |
| VM SoA | -18 | -85 | 37 | 6.710746 | 0.000001 | Cingulum_Mid_L | Left-BA31 | 1.80 | -36.94 | 32.05 | 32.83 | 36.22 | 25.41 | 435190 | Cingulum_Mid_R | Right-BA23 |
| VM BO | -39 | -46 | -20 | 6.764708 | 0.000001 | Fusiform_L | Left-Fusiform (37) | -4.76 | -20.84 | 34.05 | 33.28 | 36.34 | 22.63 | 301566 | ParaHippocampal_L | Left-Parahip (36) |
| VM BO | 30 | -49 | -17 | 4.836431 | 0.000078 | Fusiform_R | Right-Fusiform (37) | 39.80 | -56.55 | -21.17 | 12.89 | 19.41 | 13.22 | 25528 | Fusiform_R | Right-Fusiform (37) |
| VM BO | -63 | 8 | -20 | 3.934286 | 0.000708 | Temporal_Mid_L | Left-BA38 | -58.97 | -3.33 | -24.35 | 6.06 | 13.06 | 9.23 | 5906 | Temporal_Mid_L | Left-BA21 |

**Table S8 - Visuomotor whole brain correlates of the subjective experiences of Sense of Agency and Body Ownership -** MNI coordinates of clusters extracted from VM parametric maps (p<0.05, corrected) and their suggested Automated Anatomical Label and Brodmann Area (BA) output by "Label4MRI" R package.

Table S9 - Visuomotor Psychophysiological Interaction with the right insular seed

| | Peak Activation | | | | | | | Center of Gravity | | | | | | | | |
|---|---|---|---|---|---|---|---|---|---|---|---|---|---|---|---|---|
| | | | | | | | | Mean | | | STD | | | | | |
| Contrast Map | X | Y | Z | t | p | Anatomical Labeling | Brodmann Area | X | Y | Z | X | Y | Z | #Voxels | Anatomical Labeling | Brodmann Area |
| VM PPI (R Insula) | 51 | -22 | 22 | 9.377075 | p<0.00001 | Rolandic_Oper_R | Right-PrimSensory (1) | 59.02 | -24.57 | 26.32 | 4.45 | 3.68 | 6.25 | 3295 | SupraMarginal_R | Right-BA40 |
| VM PPI (R Insula) | 15 | -52 | -17 | 15.451058 | p<0.00001 | Cerebelum_4_5_R | Right-BA19 | 2.88 | -72.72 | -19.23 | 25.45 | 16.97 | 18.04 | 118177 | Vermis_6 | Right-VisualAssoc (18) |
| VM PPI (R Insula) | -3 | -1 | 1 | 22.423943 | p<0.00001 | Thalamus_L | Left-Thalamus (50) | 0.02 | -10.66 | 1.75 | 25.67 | 14.24 | 11.91 | 86820 | Thalamus_L | Right-Thalamus (50) |
| VM PPI (R Insula) | 48 | 32 | 13 | 6.943653 | p<0.00001 | Frontal_Inf_Tri_R | Right-BA46 | 46.38 | 31.39 | 13.95 | 1.58 | 1.53 | 1.51 | 160 | Frontal_Inf_Tri_R | Right-BA46 |
| VM PPI (R Insula) | 42 | 44 | 10 | 9.109797 | p<0.00001 | Frontal_Mid_R | Right-BA10 | 41.99 | 45.66 | 5.81 | 3.36 | 4.57 | 4.83 | 2630 | Frontal_Mid_R | Right-BA10 |
| VM PPI (R Insula) | 27 | 59 | -32 | -9.718437 | p<0.00001 | Frontal_Mid_Orb_R | Right-BA10 | 32.41 | 56.70 | -36.82 | 5.31 | 3.06 | 3.33 | 947 | Frontal_Mid_Orb_R | Right-BA10 |
| VM PPI (R Insula) | -3 | -7 | 52 | 11.598701 | p<0.00001 | Cingulum_Mid_L | Left-BA24 | -1.53 | -5.61 | 57.59 | 4.52 | 3.82 | 7.17 | 3505 | Supp_Motor_Area_L | Left-BA6 |
| VM PPI (R Insula) | 2 | -79 | 64 | 8.592536 | p<0.00001 | Precuneus_R | Right-BA7 | -0.73 | -76.16 | 61.30 | 3.18 | 3.33 | 2.70 | 626 | Precuneus_L | Left-BA7 |
| VM PPI (R Insula) | -9 | 47 | 22 | 9.750988 | p<0.00001 | Frontal_Sup_Medial_L | Left-BA9 | -5.64 | 42.27 | 27.25 | 4.34 | 8.38 | 12.38 | 6051 | Frontal_Sup_Medial_L | Left-BA9 |
| VM PPI (R Insula) | -9 | -25 | 46 | 7.695481 | p<0.00001 | Cingulum_Mid_L | Left-BA31 | -10.12 | -25.97 | 44.90 | 2.08 | 1.44 | 1.55 | 201 | Cingulum_Mid_L | Left-BA31 |
| VM PPI (R Insula) | -18 | 59 | 25 | 8.497411 | p<0.00001 | Frontal_Sup_L | Left-BA10 | -18.20 | 57.45 | 25.87 | 2.87 | 1.44 | 1.83 | 172 | Frontal_Sup_L | Left-BA10 |
| VM PPI (R Insula) | -20 | 65 | -38 | -9.211456 | p<0.00001 | Frontal_Sup_Orb_L | Left-BA11 | -33.61 | 57.50 | -36.38 | 6.75 | 4.50 | 3.15 | 1435 | Frontal_Inf_Orb_L | Left-BA10 |
| VM PPI (R Insula) | -24 | 41 | -8 | 7.732713 | p<0.00001 | Frontal_Mid_Orb_L | Left-BA47 | -24.90 | 41.70 | -8.07 | 2.88 | 1.45 | 0.81 | 108 | Frontal_Mid_Orb_L | Left-BA47 |
| VM PPI (R Insula) | -36 | -25 | 49 | 13.334585 | p<0.00001 | Postcentral_L | Left-PrimSensory (1) | -44.26 | -20.81 | 50.88 | 10.18 | 7.33 | 15.88 | 20027 | Postcentral_L | Left-PrimSensory (1) |
| VM PPI (R Insula) | -36 | 53 | 13 | 8.536264 | p<0.00001 | Frontal_Mid_L | Left-BA10 | -40.91 | 45.29 | 11.52 | 4.74 | 5.69 | 7.23 | 2352 | Frontal_Inf_Tri_L | Left-BA46 |
| VM PPI (R Insula) | -33 | -30 | -38 | 7.172015 | p<0.00001 | Cerebelum_6_L | Left-Parahip (36) | -32.14 | -31.00 | -36.82 | 1.29 | 0.77 | 1.51 | 57 | Cerebelum_6_L | Left-Parahip (36) |
| VM PPI (R Insula) | -36 | 29 | 10 | 7.418608 | p<0.00001 | Frontal_Inf_Tri_L | Left-BA45 | -34.62 | 29.11 | 7.74 | 1.64 | 0.89 | 3.37 | 141 | Insula_L | Left-BA45 |
| VM PPI (R Insula) | -39 | 26 | 53 | 7.352090 | p<0.00001 | Frontal_Mid_L | Left-BA8 | -40.16 | 24.60 | 50.39 | 2.29 | 1.79 | 1.76 | 261 | Frontal_Mid_L | Left-BA8 |
| VM PPI (R Insula) | -45 | -61 | 43 | 6.995578 | p<0.00001 | Angular_L | Left-BA39 | -46.92 | -59.43 | 41.09 | 2.13 | 1.60 | 1.40 | 145 | Angular_L | Left-BA39 |
| VM PPI (R Insula) | -57 | 8 | 37 | 7.291467 | p<0.00001 | Precentral_L | Left-BA6 | -54.52 | 6.66 | 37.81 | 1.87 | 4.32 | 1.93 | 339 | Precentral_L | Left-BA6 |

**Table S9 - Visuomotor Psychophysiological Interaction with the right insular seed -** MNI coordinates of clusters extracted from PPI contrast map using right insular seed (FDR corrected, q<0.05) and their suggested Automated Anatomical Label and Brodmann Area (BA) output by "Label4MRI" R package.

Table S10 - Visuomotor Psychophysiological Interaction with the left insular seed

| | Peak Activation | | | | | | | Center of Gravity | | | | | | | | |
|---|---|---|---|---|---|---|---|---|---|---|---|---|---|---|---|---|
| | | | | | | | | Mean | | | STD | | | | | |
| Contrast Map | X | Y | Z | t | p | Anatomical Labeling | Brodmann Area | X | Y | Z | X | Y | Z | #Voxels | Anatomical Labeling | Brodmann Area |
| VM PPI (L insula) | -12 | -28 | 1 | 20.109587 | p<0.00001 | Thalamus_L | Left-Thalamus (50) | -6.06 | -33.70 | -1.48 | 33.42 | 38.11 | 25.72 | 431642 | Cerebelum_4_5_L | Left-Thalamus (50) |
| VM PPI (L insula) | 27 | 59 | -31 | -10.760866 | p<0.00001 | Frontal_Mid_Orb_R | Right-BA10 | 32.66 | 57.02 | -36.86 | 5.18 | 3.21 | 3.28 | 1167 | Frontal_Mid_Orb_R | Right-BA10 |
| VM PPI (L insula) | 36 | 26 | -29 | 7.578914 | p<0.00001 | Temporal_Pole_Sup_R | Right-BA47 | 35.67 | 24.20 | -25.53 | 2.20 | 2.15 | 3.18 | 309 | Temporal_Pole_Sup_R | Right-BA47 |
| VM PPI (L insula) | 30 | -52 | 46 | 7.314073 | p<0.00001 | Parietal_Inf_R | Right-BA7 | 31.12 | -51.07 | 47.18 | 2.19 | 2.89 | 3.60 | 422 | Parietal_Inf_R | Right-BA7 |
| VM PPI (L insula) | 18 | 62 | 22 | 7.374059 | p<0.00001 | Frontal_Sup_R | Right-BA10 | 17.41 | 60.24 | 24.72 | 2.21 | 1.36 | 1.79 | 95 | Frontal_Sup_R | Right-BA10 |
| VM PPI (L insula) | -3 | 2 | 34 | 13.899351 | p<0.00001 | Cingulum_Mid_L | Left-BA24 | -2.06 | 13.20 | 40.08 | 5.72 | 21.93 | 14.87 | 38659 | Cingulum_Mid_L | Left-BA32 |
| VM PPI (L insula) | 12 | 38 | 49 | 7.596603 | p<0.00001 | Frontal_Sup_Medial_R | Right-BA8 | 12.37 | 37.80 | 50.04 | 1.49 | 1.51 | 1.42 | 127 | Frontal_Sup_R | Right-BA8 |
| VM PPI (L insula) | 3 | -79 | 63 | 8.585542 | p<0.00001 | Precuneus_R | Right-BA7 | 1.75 | -77.28 | 61.82 | 3.71 | 3.56 | 4.65 | 711 | Precuneus_R | Right-BA7 |
| VM PPI (L insula) | -3 | -37 | 28 | 6.524990 | p<0.00001 | Cingulum_Post_L | Left-BA23 | -4.42 | -37.63 | 26.94 | 1.30 | 1.20 | 1.68 | 71 | Cingulum_Post_L | Left-BA23 |
| VM PPI (L insula) | -15 | 20 | 58 | 7.021025 | p<0.00001 | Frontal_Sup_L | Left-BA6 | -15.07 | 21.00 | 58.47 | 0.63 | 1.32 | 0.99 | 30 | Frontal_Sup_L | Left-BA6 |
| VM PPI (L insula) | -15 | 44 | 40 | 7.201221 | p<0.00001 | Frontal_Sup_L | Left-BA8 | -17.30 | 42.08 | 39.36 | 2.01 | 1.22 | 2.19 | 115 | Frontal_Sup_L | Left-BA8 |
| VM PPI (L insula) | -21 | 35 | 58 | 7.981019 | p<0.00001 | Frontal_Sup_L | Left-BA8 | -19.46 | 32.84 | 57.52 | 2.33 | 1.78 | 1.59 | 231 | Frontal_Sup_L | Left-BA8 |
| VM PPI (L insula) | -21 | -73 | 43 | 7.291690 | p<0.00001 | Parietal_Sup_L | Left-BA7 | -19.63 | -73.02 | 43.74 | 1.95 | 1.97 | 1.95 | 317 | Parietal_Sup_L | Left-BA7 |
| VM PPI (L insula) | -18 | -67 | 34 | 7.831288 | p<0.00001 | Precuneus_L | Left-BA7 | -18.94 | -66.11 | 34.23 | 1.16 | 1.42 | 1.72 | 111 | Occipital_Sup_L | Left-BA7 |
| VM PPI (L insula) | -24 | 41 | -8 | 7.596076 | p<0.00001 | Frontal_Mid_Orb_L | Left-BA47 | -24.79 | 40.53 | -9.07 | 3.50 | 2.43 | 1.27 | 276 | Frontal_Mid_Orb_L | Left-BA47 |
| VM PPI (L insula) | -24 | 65 | -34 | -9.677835 | p<0.00001 | Frontal_Sup_Orb_L | Left-BA10 | -33.33 | 57.61 | -36.24 | 6.73 | 4.47 | 3.12 | 1631 | Frontal_Inf_Orb_L | Left-BA10 |
| VM PPI (L insula) | -27 | -67 | 55 | 6.746190 | p<0.00001 | Parietal_Sup_L | Left-BA7 | -26.50 | -66.75 | 54.46 | 0.87 | 1.18 | 0.68 | 28 | Parietal_Sup_L | Left-BA7 |
| VM PPI (L insula) | -39 | 17 | 28 | 6.921340 | p<0.00001 | Frontal_Inf_Tri_L | Left-BA44 | -38.58 | 17.32 | 28.77 | 1.71 | 1.48 | 1.68 | 159 | Frontal_Inf_Tri_L | Left-BA44 |
| VM PPI (L insula) | -45 | 27 | -32 | 7.451139 | p<0.00001 | Temporal_Pole_Sup_L | Left-BA38 | -46.16 | 22.88 | -32.60 | 1.63 | 1.96 | 2.25 | 253 | Temporal_Pole_Mid_L | Left-BA38 |
| VM PPI (L insula) | -57 | 14 | -32 | 7.896594 | p<0.00001 | Temporal_Pole_Mid_L | Left-BA38 | -56.29 | 10.08 | -32.90 | 1.13 | 2.18 | 1.64 | 133 | Temporal_Pole_Mid_L | Left-BA38 |
| VM PPI (L insula) | -60 | -25 | -14 | 7.116969 | p<0.00001 | Temporal_Mid_L | Left-BA21 | -60.14 | -25.24 | -14.47 | 1.58 | 0.87 | 1.05 | 59 | Temporal_Mid_L | Left-BA21 |

**Table S10 - Visuomotor Psychophysiological Interaction with left insular seed -** MNI coordinates of clusters extracted from PPI contrast map using left insular seed (FDR corrected, q<0.05) and their suggested Automated Anatomical Label and Brodmann Area (BA) output by “Label4MRI” R package.

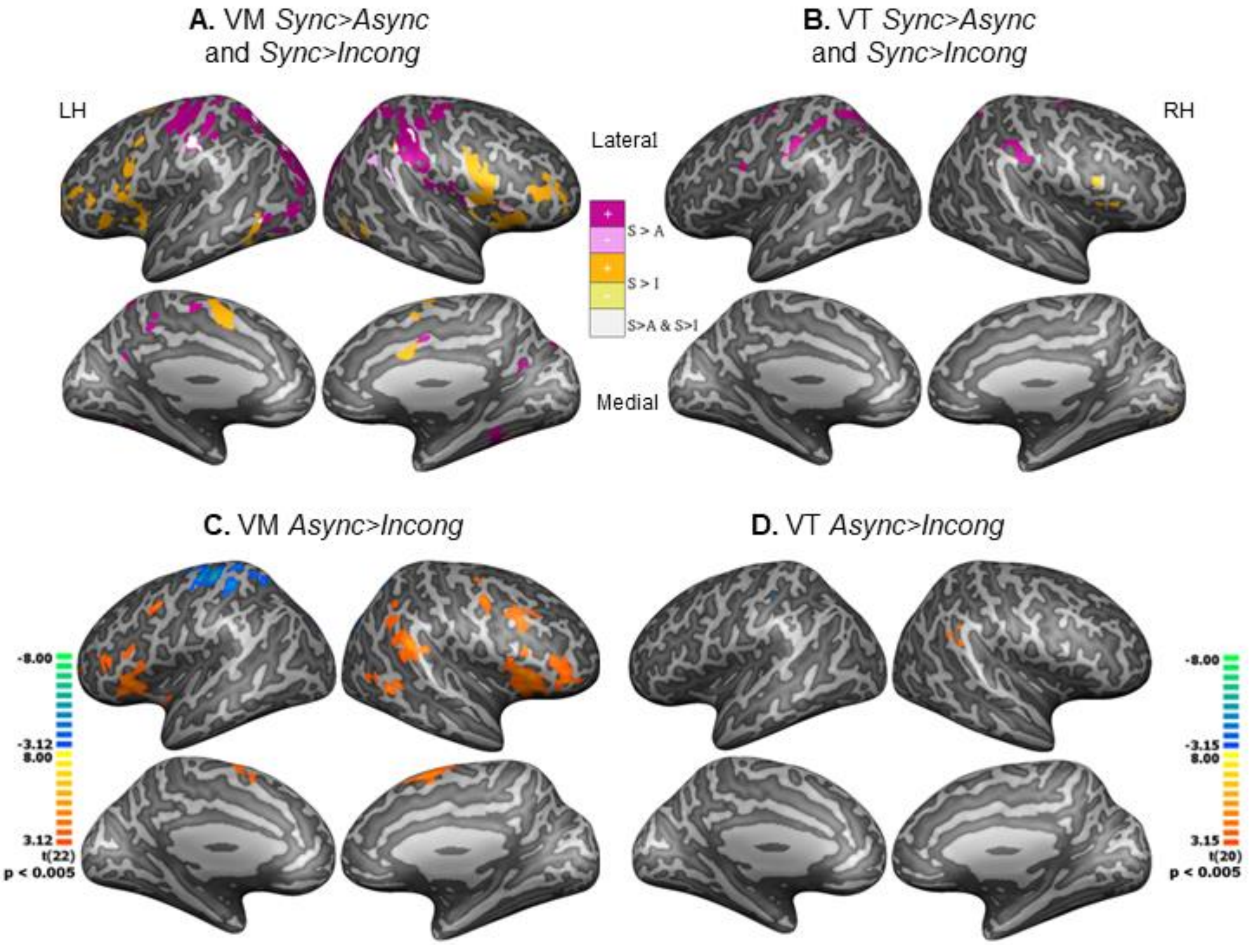


**Figure S1 - Divergent neural processing for temporal and anatomical conflicts -** **(A)** VM *Sync>Async* (temporal conflict) and *Sync>Incong* (anatomical conflict) contrast maps overlaid and displayed in solid colors, purple and orange clusters were more activated during *Sync* condition while pink and yellow were more activated during *Async* and *Incong* conditions respectively, overlapping clusters are displayed in white. (**B)** VT *S > A* and *S > I* contrast maps overlaid. **(C)** VM *Async>Incong* contrast map (p = 0.005, cluster threshold: 55 voxels). **(D)** Visuotactile (VT) *Async>Incong* contrast map (p = 0.005, cluster threshold: 38 voxels). All maps are displayed in neurological orientation, thresholded at p<0.005, and corrected for multiple comparisons with a cluster-size threshold.

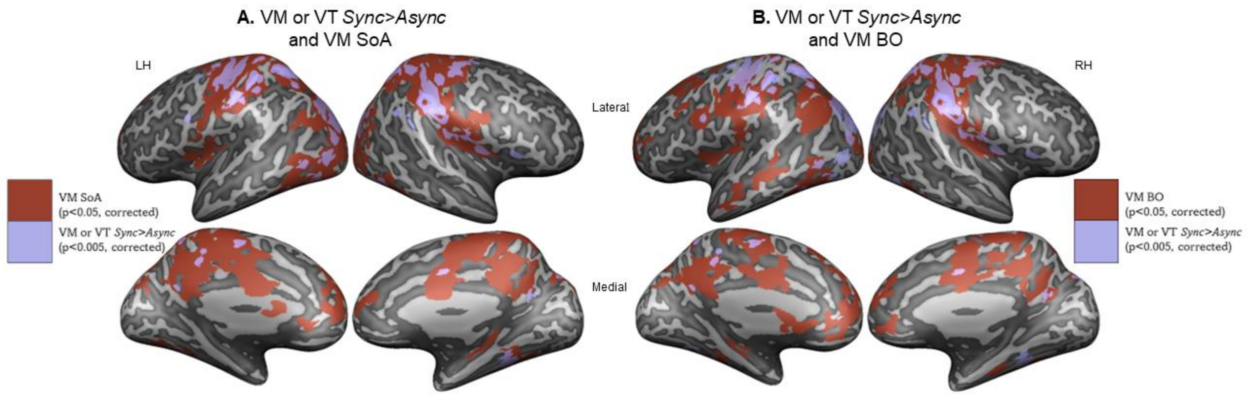


**Figure S2 - Dissociation of sensory stimulation and subjective experience - (A)** VM SoA (red) and the union (or) of VM and VT *Sync>Async* (light purple) **(B)** VM BO and the union of VM and VT *Sync>Async*. Note: left Insula, Anterior Cingulate Cortex (ACC), ventromedial Prefrontal Cortex (vmPFC), and precuneus activity followed the subjective experience of BO and SoA but were not active during illusory stimulation.

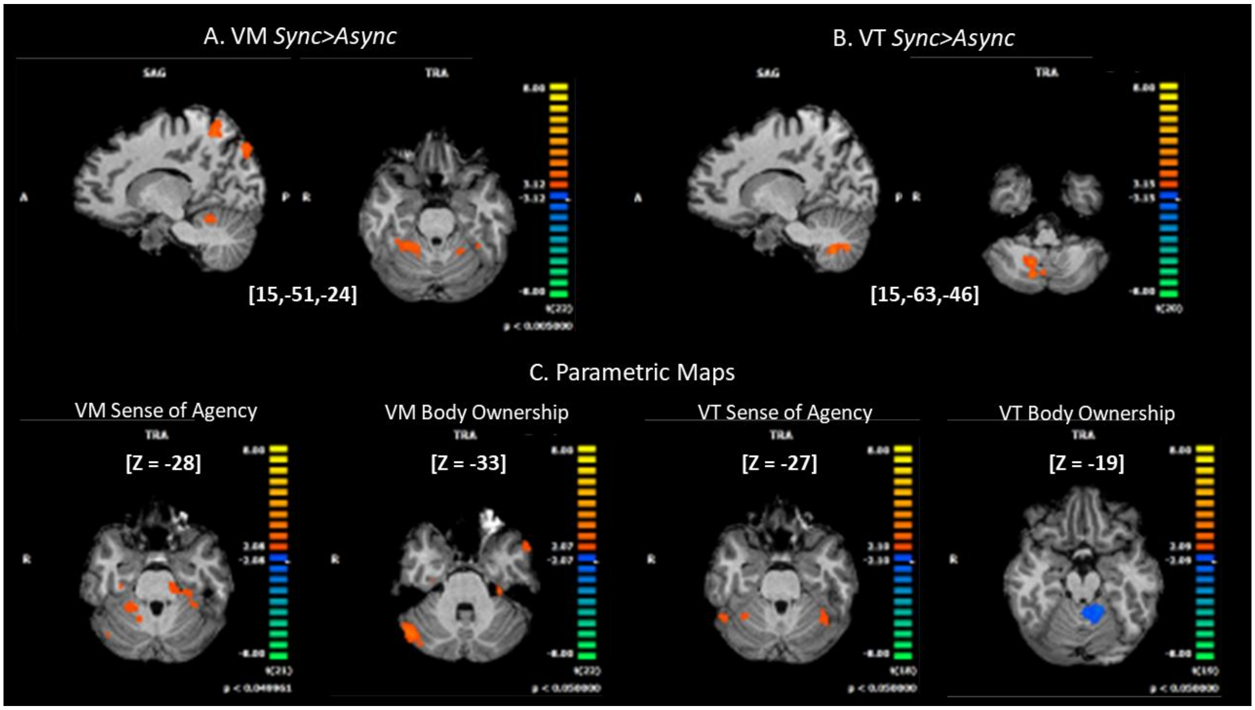


**Figure S3 - Cerebellar activity during illusion relates to Agency and Ownership - (A)** VM and **(B)** VT *Sync>Async* contrast maps showing significant cerebellar activity during the illusion. **(C)** Cerebellar correlates of SoA and BO, all slices are in radiological view.
